# Distinct roles for two *Caenorhabditis elegans* acid-sensing ion channels in an ultradian clock

**DOI:** 10.1101/2021.11.15.468673

**Authors:** Eva Kaulich, Brian D. Ackley, Yi-Quan Tang, Iris Hardege, William R. Schafer, Denise S. Walker

## Abstract

Biological clocks are fundamental to an organism’s health, controlling periodicity of behavior and metabolism. Here, we identify two acid-sensing ion channels, with very different proton sensing properties, and describe their role in an ultradian clock, the defecation motor program (DMP) of the nematode *Caenorhabditis elegans*. An ACD-5-containing channel, on the apical membrane of the intestinal epithelium, is essential for maintenance of luminal acidity, and thus the rhythmic oscillations in lumen pH. In contrast, the second channel, composed of FLR-1, ACD-3 and/or DEL-5, located on the basolateral membrane, controls the intracellular Ca^2+^ wave and forms a core component of the master oscillator that controls timing and rhythmicity of the DMP. *flr-1* and *acd-3/del-5* mutants show severe developmental and metabolic defects. We thus directly link the proton-sensing properties of these channels to their physiological roles in pH regulation and Ca^2+^ signaling, the generation of an ultradian oscillator, and its metabolic consequences.

**One-Sentence Summary:** Two acid-sensing DEG/ENaC channels play distinct roles in controlling different aspects of rhythmic proton and Ca^2+^ oscillations in the *C. elegans* intestine.

## Introduction

Many physiological processes occur with a predictable periodicity. The maintenance of such rhythms can rely on fluctuating gene expression, hormone concentrations, or homeostatic oscillations in signaling molecules within cellular compartments, depending on the timescale of the clock (*1–4*). Disruption of biological clocks can have devastating consequences on homeostasis or behavior (*5*). Here we examine the physiological role of acid-sensing ion channels (ASICs) in the defecation motor program (DMP) of the nematode *Caenorhabditis elegans*, an ultradian rhythm that occurs about every 50 seconds (*3, 4*). We take advantage of this short-period cellular oscillator to relate ion channel physiology *in vivo* to organismal metabolism and behavior.

ASICs, the acid-sensing members of the Degenerin/Epithelial Sodium Channel (DEG/ENaC) superfamily of cation channels are thought to be the main proton receptors in vertebrates (*6, 7*). They are an important drug target, because acidosis is a feature of painful inflammatory and ischemic conditions. ASICs are widely expressed in the central and peripheral nervous system, where their roles in inflammation, ischemia, pain perception, and learning are extensively studied (*8–13*). However, less is known about their physiological role in non-neuronal tissue such as the intestinal epithelium, despite cross-phyla evidence for gastrointestinal expression and the role of acidity therein (*14–17*). The extreme acidity of the stomach requires an elaborate system of mucosal protective mechanisms, failure of which results in conditions such as gastritis, ulceration and dyspepsia. Acidification of the intestinal tract is also a hallmark of conditions such as irritable bowel syndrome and cystic fibrosis, so understanding the role of acid sensors, of which the ASICs are prime candidates, is of wide-ranging therapeutic importance (*18*).

The DMP depends on inositol 1,4,5-trisphosphate (IP_3_)-dependent Ca^2+^ oscillations in the intestinal epithelial cells. IP_3_ signaling loss-of-function mutations cause dramatic increases in both period length and variability, whereas overexpression of the IP_3_ receptor, ITR-1, reduces period length (*19–21*). Increased cytosolic Ca^2+^ results in the rhythmic release of protons from the basolateral membrane, into the pseudocoelom. This triggers the posterior body wall muscles to contract (pBoc), via activation of a proton-gated cation channel PBO-5/PBO-6 (*22, 23*). Propagation of the Ca^2+^ wave from the posterior to anterior then initiates contraction of the body wall muscle at the anterior of the intestine (aBoc) a few seconds later, via activation of the motor neuron AVL, and then finally, GABA release from both AVL and DVB triggers the enteric muscle contraction (EMC), expelling gut contents (*24, 25*). Premature termination of the Ca^2+^ wave or pharmacologically blocking the Ca^2+^ wave propagation before it can reach the anterior disrupts aBoc and EMC (*26*), further supporting the idea that Ca^2+^ is the master regulator.

In addition to Ca^2+^ and pseudocoelomic pH, proton oscillations in the intestinal lumen are also integral to the DMP. The luminal pH fluctuates between ∼4.4 and 6.5 during the cycle and the pH transitions in a wave that travels from posterior to anterior (*22, 23, 27-29*), dependent on proton uptake and release at the apical membrane. Proton gradients are fundamental to nutrient absorption, and some DMP-affecting mutations can result in aberrant fat metabolism and deficient nutrient uptake (*27, 30*). Loss of proton regulation can also disrupt immunity and pathogen resistance, as seen in mutants for *C. elegans* CHP1 homolog, PBO-1, in which the lumen is constantly acidic (*31*). Thus, dysregulation of components of this ultradian clock can have far reaching organismal consequences.

The precise relationship between luminal proton oscillations, intestinal Ca^2+^ and the DMP is unclear; but the mutant phenotypes of some of the known transporters and exchangers that control lumen pH provide some clues. Knock-down of VHA-6, a subunit of the apical proton-transporting V-type ATPase, leads to more neutral lumen pH and an increase in cycle length (*27*). Mutations in the Na^+^/H^+^ exchanger PBO-4 also cause a more neutral lumen pH, but cycle length and Ca^2+^ oscillations are unaffected (*4, 22, 23, 28, 31*). Mutants of *pbo-1*, on the other hand, have a highly acidic lumen and no proton wave, but Ca^2+^ oscillations and cycle length are largely unaffected (*4, 23, 31, 32*). This suggests that periodicity of the DMP can be independent of luminal pH, and that proton dynamics are likely downstream of the Ca^2+^ oscillator.

We hypothesized that a proton wave transitioning from the posterior to the anterior of the lumen could lead to sequential activation of proton sensors in the apical membrane (*28, 29, 31*), and acid-sensing DEG/ENaCs are good candidates. Compared to mammals, the *C. elegans* genome encodes a vastly expanded array of 30 DEG/ENaCs. One of these, FLR-1 (FLuoride-Resistant phenotype), functions in DMP rhythmicity and execution (*16, 33*), although, the mechanistic basis of this role is unclear. With the exception of ACD-1, the first acid-inactivated DEG/ENaC to be identified (*34*), the proton sensing properties of the *C. elegans* DEG/ENaCs have not been characterized.

Here, we address the physiological role of *C. elegans* acid-sensing DEG/ENaCs in the intestine. We explore how Ca^2+^ and proton oscillations maintain periodicity and execution of the DMP, and find that disruption of Ca^2+^ and proton homeostasis are detrimental to metabolism and organismal health. We show that four intestinal DEG/ENaC subunits form two acid-inactivated channels with distinct properties. The first contains ACD-5 and localizes to the luminal face of the intestinal cells and acts as a proton sensor, establishing low luminal pH and thus enabling cyclic pH oscillations. The second channel, containing FLR-1, ACD-3 and/or DEL-5, localizes to the basolateral membrane, facing the pseudocoelom, and is essential for the Ca^2+^ oscillations that control the timing and execution of the DMP and thus is a key component of this ultradian clock. We thus provide direct evidence linking the channel properties of these proton sensors to their roles in pH regulation and Ca^2+^ signaling *in vivo*, which can provide insight into potential cellular roles for other acid-sensing members of this important channel family.

## Results

### ACD-5 forms a homomeric proton-sensing channel localized to the apical membrane

Acid-sensing members of the DEG/ENaC family can form homomeric or heteromeric channels that are activated or inhibited by low pH (*34–36*). We expressed 24 *C. elegans* DEG/ENaC subunits in *Xenopus* oocytes and assayed for activity using Two-Electrode-Voltage clamp (TEVC) recording. Oocytes expressing the ACD-5 subunit showed currents that could be inhibited by amiloride, a known DEG/ENaC blocker, (Figure 1A, B), and showed robust responses to pH. ACD-5 exhibited maximal current at around pH 6 and was inhibited by both higher and lower pH, with pH_50_s of 4.87 and 6.48 (Figure 1C).

**Figure 1.**
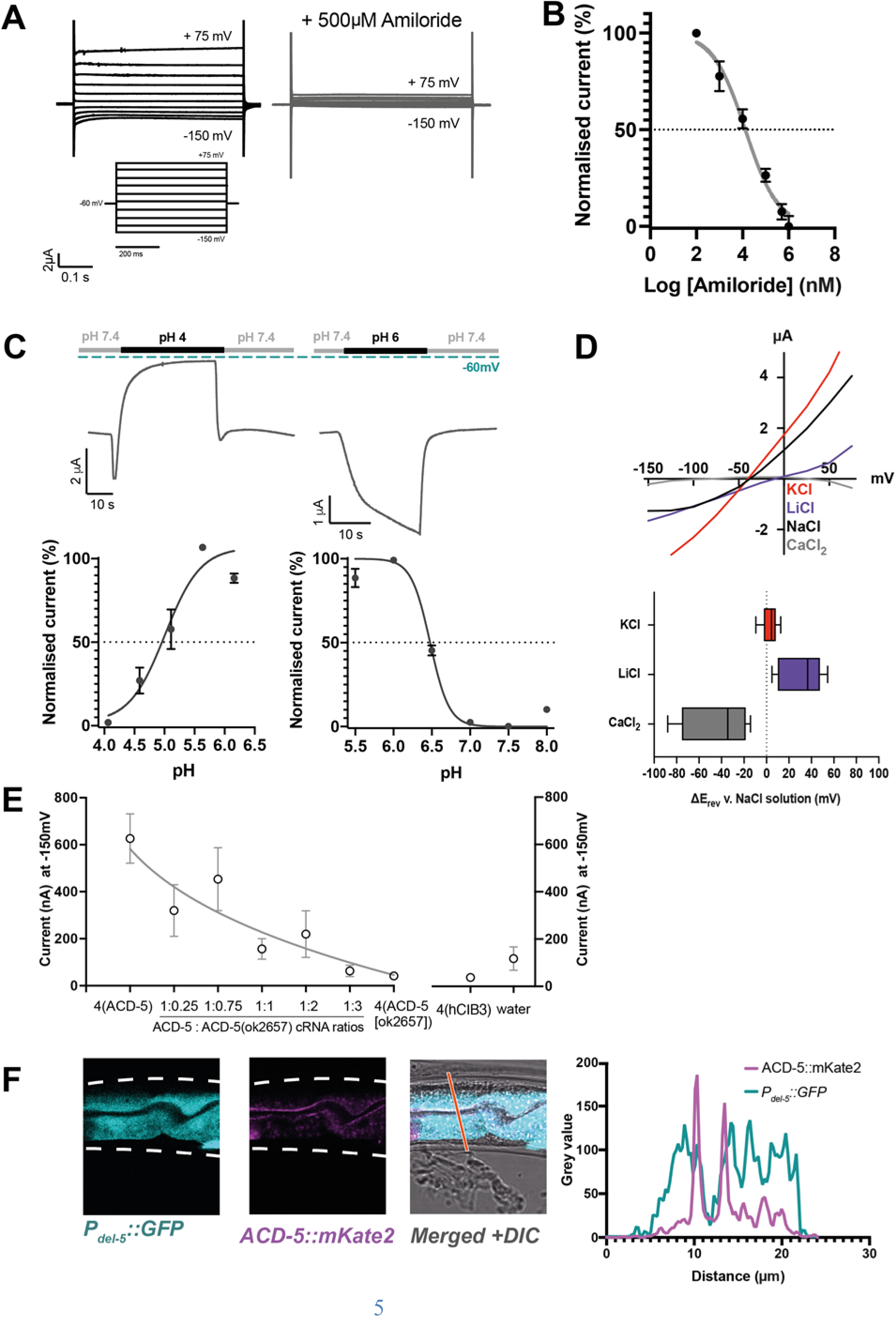
ACD-5 is an amiloride-sensitive, acid-sensing cation channel on the luminal membrane of the intestine. (**A**) ACD-5 currents are amiloride sensitive. Representative ACD-5 transients in *Xenopus* oocytes in the presence and absence of amiloride and (**B**) Normalized (I/I_max_) amiloride-dose response curve for ACD-5, showing a half maximal inhibitory concentration (IC_50_) of 131 μM (LogIC_50_ = 4.118) (N = 8) indicated by the dashed line. (Curve fitted as described below). Error bars represent Mean ± SEM. (**C**) ACD-5 can form a homomeric acid-sensing ion channel. Upper panel: Representative trace of ACD-5 injected *Xenopus* oocytes subjected to a down-step to pH 4 or 6. Currents were recorded at a holding potential of -60mV and traces are baseline-subtracted and drift-corrected using the Roboocyte2+ (Multichannels) software. Lower panel: pH-dose dependence. ACD-5-epxressing oocytes were perfused with solutions of decreasing pH from pH 6 (left), showing an inhibitory pH_50_ of 4.90 (N = 5); when perfused with increasing pH from pH 6 (right), showing an excitatory pH_50_ of 6.48 (N = 7). Currents were recorded at a holding potential of -60mV, normalized to maximal currents and best fitted with the Hill’s equation (Nonlin fit Log(inhibitor) vs normalized response – variable slope) in GraphPad Prism. Error bars represent Mean ± SEM. (**D**) Current-voltage (IV) curve (left) and change in reversal potential, ΔE_rev_ when shifting from a NaCl solution to CaCl_2_, KCl or LiCl. N = 12 oocytes. Presented as Median and IQR (Tukey method). (**E**) ACD-5(ok2657) is a dominant mutation. Expression of varying ratios of ACD-5(ok2657) with a constant concentration of wild-type ACD-5 in *Xenopus* oocytes. hCIB3 (Calcium and integrin-binding family member 3) was used as a filler to account for amount of cRNA injected. Mean ± SEM. 9<N>19 for each ratio. (**F**) ACD-5 is localized at the apical (luminal) intestinal membrane. Localization of mKate2-C-terminally tagged ACD-5 (*ljEx1470 (Pacd-5::acd-5(no stop) cDNA::mKate2))* shows apical membrane localization (magenta). Cytoplasm of the intestinal cells is shown in cyan (*ljEx1349 (Pdel-5::GFP)*). Intensity profile taken at the part of the intestine indicated by the red line. Scale bar:10 μm.

We next investigated ion selectivity, assessing the shift in reversal potential (ΔE_rev_) by substituting NaCl with either equimolar KCl, LiCl or a CaCl_2_ (*34, 37*). Monovalent ion-substitution experiments were conducted in the absence of Ca^2+^, as Ca^2+^ can block some vertebrate ASICs (*38*). Permeability for Ca^2+^ (*34*) was also tested, as some constitutively open *C. elegans* DEG/ENaCs are permeable for Ca^2+^ in addition to monovalent cations (*39*). ACD-5 shows a preference for Li^+^ over Na^+^ and a small preference of K^+^ over Na^+^ and is Ca^2+^ impermeable (Figure 1D). ACD-5 was not permeable for protons, as the ΔE_rev_ did not show a positive shift when extracellular proton concentration was increased, and removal of Ca^2+^ from the solution also did not alter ΔE_rev_. By contrast, removal of Na^+^ (by substituting with NMDG) induced a large negative ΔE_rev_ (Figure 1-figure supplement 1A-D). Thus, ACD-5 forms a homomeric amiloride-sensitive, pH-sensitive monovalent cation channel.

To determine the function of the ACD-5 channel, we investigated its mutant phenotype. The only *acd-5* mutation available was *acd-5(ok2657),* generated by the *C. elegans* Deletion Mutant Consortium (*40*), a truncation lacking TMD2 and the C-terminus (Figure 1-figure supplement 2A, B). To determine the effect of this mutation on channel function, we expressed the mutant form in *Xenopus* oocytes. Oocytes expressing ACD-5(ok2657) alone did not display any currents and were unaffected by amiloride (Figure 1-figure supplement 3A, B). However, co-expressing ACD-5(ok2657) with wild-type ACD-5 significantly altered the expressed currents. Specifically, increasing the ratio of injected *acd-5(ok2657)* cRNA to wild-type *acd-5* cRNA resulted in a decrease in current (Figure 1E). This suggests that the ACD-5(ok2657) subunit either forms a non-functional heteromer or sequesters factors required for wild-type ACD-5 function, and thus is a dominant mutation, hindering functional channel formation.

Given ACD-5’s dual response to protons, we investigated its subcellular localization to understand its physiological pH environment. A translational *acd-5::mKate2* C-terminal fusion showed expression in the intestinal epithelial cells, localizing to the apical membrane, as confirmed by co-localization with an apical membrane marker (OPT-2::GFP (*41*)) and by comparing the localization with that of markers for the intestinal cytoplasm (*Pdel-5::GFP*) and lumen (ingested fluorescent dye, DiO) (Figure 1F and Figure 1-figure supplement 4). The observation that ACD-5 localizes to the apical membrane indicated that its physiological environment is likely in the range pH 4-6, the pH range of the intestinal lumen (*23, 29*), consistent with our electrophysiological findings that ACD-5 exhibits maximal current at pH 6 and is inhibited at pH 4.

### Mutations in acd-5 dramatically alter intestinal lumen pH

Given its luminal location and physiological sensitivity to mildly acidic pH, we hypothesized ACD-5 may have an impact on the acidity of the lumen. To test this, we used the ingested pH-sensitive fluorescent dye Kansas Red (KR35) that allows visualization of the dynamic wave in luminal pH. Every 50 seconds, the posterior fluorescence transitions to the anterior region of the intestine, remaining there for 3-7 seconds, a period referred to as the Maximum Anterior Transition (MAT) (*28, 31*). We analyzed both the dominant *acd-5(ok2657)* as well as a null mutant generated using CRISPR/Cas9 (*42*), *acd-5(lj122)* (Figure 1-figure supplement 2A, B). Mutants carrying either *acd-5* allele showed overall lower fluorescence intensities, indicating a more neutral lumen pH compared to wild-type (Figure 2A, B). As a result, both *acd-5(lj122)* and *acd-5(ok2657)* mutants exhibited a high incidence of missed (or undetectable) MATs, with only 61 % and 56 %, respectively, displaying at least one MAT in the 2-minute recording time, compared with 82 % of wild-type animals (Figure 2C). These defects were rescued in the *acd-5(lj122)* null by expression of the wild-type *acd-5* gene, but rescue was not observed for the truncated *acd-5(ok2657)* (Figure 2A, B). Where detectable, MAT interval length was not grossly affected (Figure 2D). Thus, although ACD-5 apparently plays a pivotal role in the control of proton concentration, it does not greatly influence the timing of the luminal proton wave.

**Figure 2.**
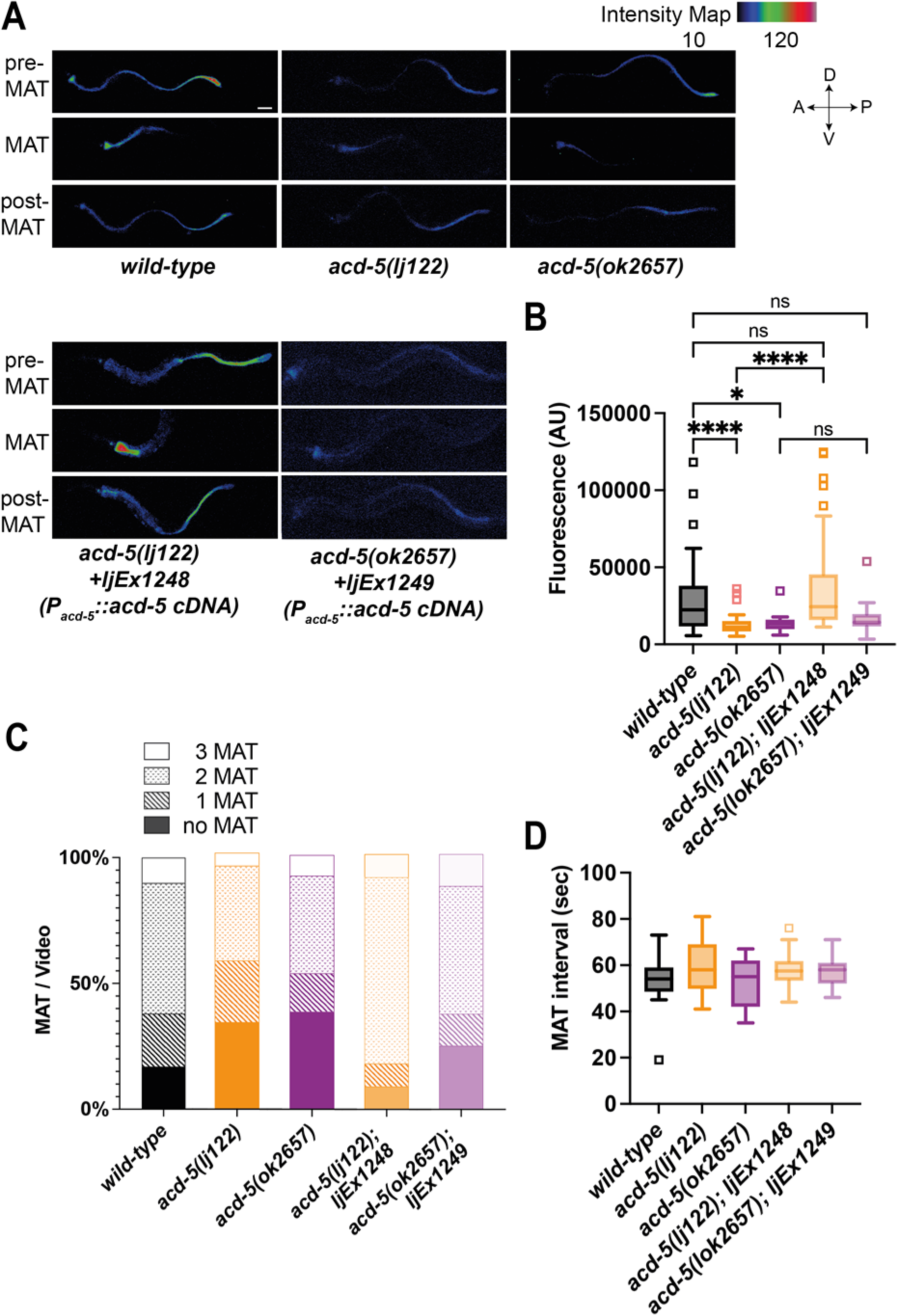
ACD-5 mutations show disruptions in intestinal lumen acidity and luminal proton oscillations. (**A**) ACD-5 is important to establish and maintain an acidic intestinal pH. Heat map of pixel fluorescence intensity with red representing highest intensity (high acidity), and black lowest intensity (low acidity), after feeding on the pH-sensitive probe Kansas Red (KR35) for 30 min (10 μM). Maximum anterior transition (MAT) ± 15 seconds (pre/post-MAT) during the DMP extracted from videos of free moving animals are shown. N = 24, 19, 9, 20, 12, respectively. Rescue lines express the *ljEx1248(Pacd-5::acd-5 cDNA)* or *ljEx1249(Pacd-5::acd-5 cDNA)* extrachromosomal array, respectively. D=dorsal, V=ventral, A=anterior, P=posterior. Scale bar: 50 μm. (**B**) *acd-5(lj122)* mutants (\*\*\*\**p*<0.0001) and *acd-5(ok2657)* mutants (\**p*=0.0309) display lower intestinal lumen pH compared to the wild-type. This could be rescued only in the *acd-5(lj122)* mutants (\*\*\*\**p*<0.0001). Quantification of fluorescence in the MAT. Data is presented as median and IQR (Tukey method). Outliers are indicated by squared boxes. Kruskal Wallis test with Dunn’s multiple comparison test. (**C**) *acd-5* mutants display a reduced number of MATs. Quantification of MATs per 2-minute video recording shown as percentages of animals displaying 1, 2 or three MATs. N = 29, 38, 13, 23, 16, respectively. (**D**) MAT intervals are not affected by *acd-5* mutations. Time interval of MAT of *acd-5* mutants, wild-type and rescues. Kruskal Wallis test with Dunn’s multiple comparison test was non-significant. Data is presented as median and IQR (Tukey method). N = 21, 18, 7, 20, 12, respectively.

### Mutations in acd-5 subtly alter intestinal Ca^2+^ oscillations and DMP behavior

Mutations that disrupt proton homeostasis in the intestinal lumen can have differing effects on intestinal Ca^2+^ transients and DMP behavioral output. Therefore, we assessed whether *acd-5* affects intestinal Ca^2+^ oscillations, the presumed master oscillator for the DMP. Both alleles displayed intact, regular Ca^2+^ oscillations, with small changes in timing and amplitude. *acd-5(ok2657)* mutants showed an increased amplitude and significantly longer intervals between Ca^2+^ peaks, whereas *acd-5(lj122)* null animals displayed a reduced amplitude and a very small decrease in interval length (Figure 3A, B and Figure 3-figure supplement 1 for representative example traces). This shows that *acd-5* does interact with the Ca^2+^ oscillations, albeit in a relatively subtle manner.

**Figure 3.**
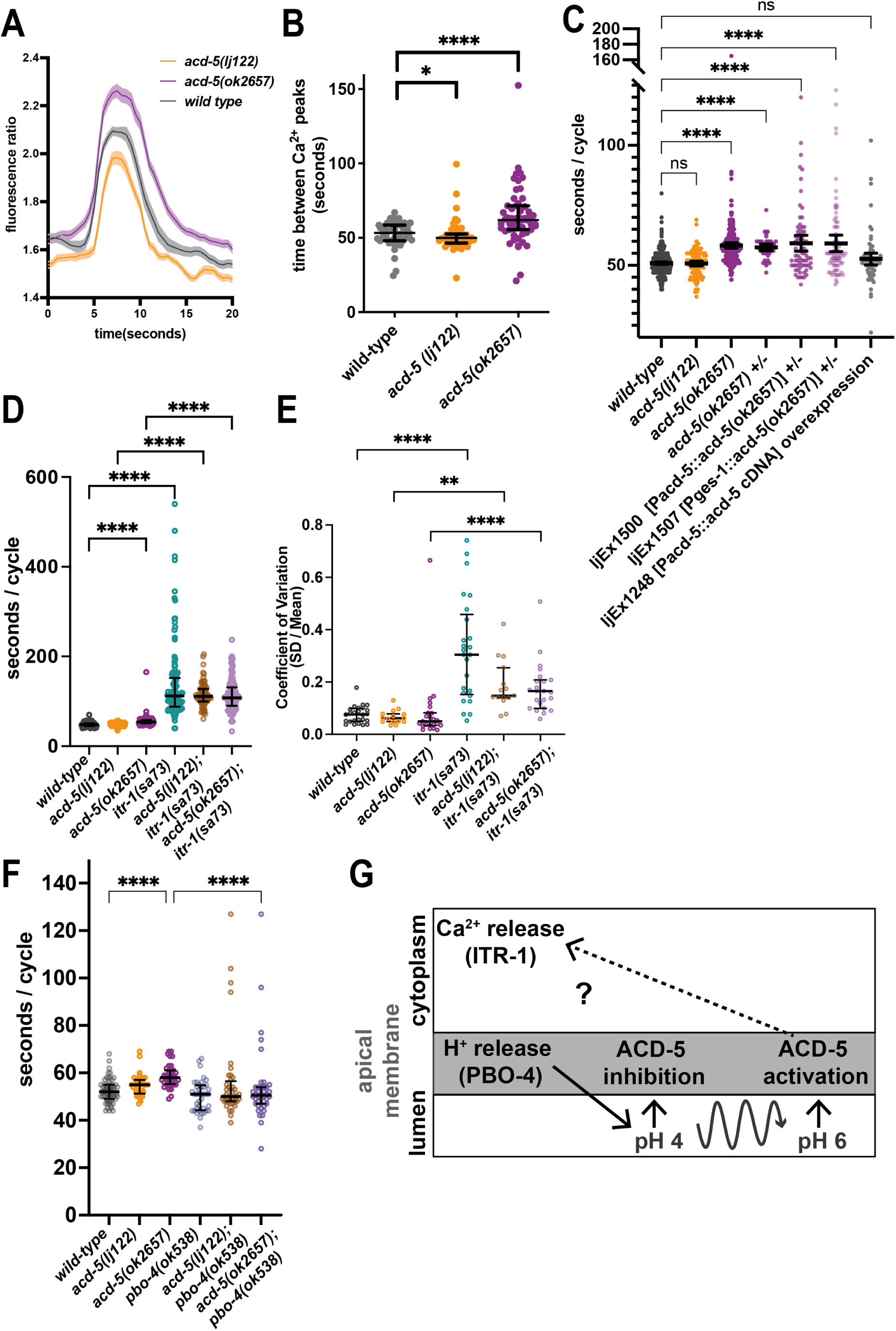
ACD-5 mutations affect Ca^2+^ oscillations and behavior. (**A**) *acd-5* mutations affect intestinal Ca^2+^ oscillations. Ca^2+^ mean trace ± SEM of fluorescence ratio for *acd-5* mutants and wild-type. (**B**) Ca^2+^ cycle length. Mann-Whitney U between wild-type and *acd-5(ok2657)* \*\*\*\**p*<0.0001 (indicating a longer cycle interval), and between wild-type and *acd-5(lj22) *p*= 0.0266 (indicating a shorter cycle interval). Error bars represent median and IQR, N =50, 55 and 53, respectively (i.e. individual intervals from 15 animals for each genotype, recorded for 5 min each). (**C**) ACD-5(ok2657) is a dominant mutation. Comparison of defecation cycle length in *acd-5* mutant and overexpression strains. Kruskal-Wallis test and post-hoc Dunnett’s multiple comparisons test found no significant difference between wild-type and *acd-5(lj122)* animals (p>0.9999) but a significant difference from wild type for homo- and heterozygous *acd-5(ok2657)* animals and wild-type animals expressing *ljEx1500(Pacd-5::acd-5[ok2657])* or *ljEx1507(Pges-1::acd-5[ok2657])* (****p<0.0001); those overexpressing wild-type *ljEx1248(Pacd-5::acd-5cDNA)* were not significantly different from wild type (p>0.9999). Median and IQR. 50>N>75 animals (10-15 animals for each group, 5 cycles for each animal). (**D**) Effect of *acd-5* alleles on the defecation interval of *itr-1(sa73)* animals. DMP interval length (median and IQR). Kruskal-Wallis test was significant (\*\*\*\**p*<0.0001). A post-hoc Dunn’s multiple comparison test indicated that compared to the wild-type, *acd-5(ok2657)* mutants and the *itr-1* single and double mutants (\*\*\*\**p*<0.0001) all showed a significant increase in interval length. N = 45 (15 animals x 5 cycles for each). (**E**) Coefficient of variation in cycle length for *itr-1* and *acd-5* single and double mutants. *acd-5(lj122)* (\*\**p*=0.001) or the *acd-5(ok2657)* (\*\*\*\**p*<0.0001) double mutants showed significant increase in variability, similar to the *itr-1* single mutants. Error bars represent median and IQR. (15 animals x 5 cycles for each). (**F**) Effect of *acd-5* alleles on defecation cycle length of *pbo-4(ok538)* animals (N= 12 for the wild type, 8 for each mutant x 5 cycles each). DMP interval length (median and IQR). The Kruskal-Wallis test was significant with a post-hoc Dunn’s multiple comparison test indicating that only *acd-5(ok2657)* mutants had increased cycle length (\*\*\*\**p*<0.0001); *pbo-4* single and double mutants were not significantly different from wild-type. *pbo-4(ok538)* significantly reduced the cycle length of *acd-5(ok2657)* animals (\*\*\*\**p*<0.0001). Error bars represent median and IQR. (**G**) Schematic of the relationship between ACD-5, proton release and Ca^2+^ oscillations. Release of protons at the luminal membrane via PBO-4 results in decrease in luminal pH which inhibits the ACD-5 channel, proton oscillations (curved line) form pH 4 to pH 6 activate the ACD-5 channel at pH 6 which in turn influences (but does not drive) *itr-1*-mediated Ca^2+^ oscillations.

To explore whether the changes observed in Ca^2+^ oscillation translate into behavioral outputs, we assessed the timing and execution of DMP cycles. The cycle intervals observed correspond well with the Ca^2+^ imaging observations; *acd-5(ok2657)* exhibited significantly longer intervals and *acd-5(lj122)* animals showed no difference from wild type (Figure 3C). The distinct phenotypes observed for these two alleles were consistent with our *in vitro* evidence that *acd-5(ok2657)* is a dominant mutation. Heterozygous animals and wild type animals overexpressing the *acd-5(ok2657)* mutant gene, under the endogenous promoter or under another intestinal promoter (*Pges-1*) all retained the longer cycle phenotype, supporting this hypothesis (Figure 3C). We also found that overexpression of the *acd-5(ok2657)* gene subtly increased variability and disrupted EMCs, although since this was also true of overexpression of the wild-type *acd-5* gene (Figure 3-figure supplement 2A,B), it may have been a consequence of overexpression rather than the *acd-5(ok2657)* mutation itself.

To gain further insight into the role of *acd-5* in the DMP, we examined the genetic interaction between *acd-5* alleles and known DMP factors. The IP_3_ receptor, ITR-1, plays a critical role in the Ca^2+^ oscillations that drive the DMP and he loss-of-function allele *itr-1(sa73)* results in a dramatic increase in both cycle length and variability (*19–21*). We observed the previously described compensatory effect (*33*) in *itr-1(sa73),* where an aberrantly long cycle is followed by a short one, and vice versa (Figure 3-figure supplement 3A). Although the *acd-5* alleles did not significantly reduce the cycle length or variability of *itr-1(sa73)*, and the compensatory effect remained intact, we did nevertheless observe a striking suppression of excessively long cycles, with an almost complete loss of cycles over 200 seconds (Figure 3D, E and Figure 3-figure supplement 3A-C). This suggests that *acd-5* is responsible for the dysregulation that results in those longer cycles, perhaps suggesting a role for ACD-5 in regulating cycle length that is otherwise presumably masked or overridden by the dominant influence of the Ca^2+^ oscillator.

Mutations in the Na^+^/H^+^ exchanger gene, *pbo-4* show a more neutral luminal pH, with fewer MATs observed during the proton wave, similar to *acd-5* mutants (*31*). NHX-mediated luminal pH increases immediately preceding the peak of the Ca^2+^ wave and pBoc (*27*), however, the precise interplay between *itr-1*-mediated Ca^2+^ increase and *pbo-4*-mediated proton oscillations remains unknow. PBO-4 also localizes to the basolateral membrane and mutations result in missed or weak pBoc and frequently missed EMCs (*22, 31*). This dual role complicated our interpretation of its interaction with *acd-5*. The *acd-5/pbo-4* double mutants both displayed wild-type cycle length and variability, and EMC and pBoc defects similar to the *pbo-4* mutant (Figure 3F, Figure 3-figure supplement 3D-F). This is consistent with *pbo-4* acting genetically upstream of *acd-5* and lack of effect on pBoc is consistent with the apical role of *acd-5* (Figure 3G).

### FLR-1, ACD-3 and DEL-5 localize to the basolateral membrane and can form acid-sensitive heteromeric channels

Previous research identified expression of another DEG/ENaC channel gene, *flr-1,* in the intestine (*16*), and it was shown to localize to both apical and basolateral membranes (*16, 43*). *flr-1* mutants show more severe disruption of the DMP than in *acd-5*, leading us to hypothesize that there might be two distinct channels in the intestinal cell: A heteromeric or homomeric ACD-5-containing channel on the apical membrane and a FLR-1-containing channel on both membranes. Since FLR-1 expressed alone in oocytes did not produce pH sensitive currents in our experiments, we hypothesized that FLR-1 could contribute to a heteromeric channel. Therefore, we used transcriptional fusions to search for additional intestinally expressed DEG/ENaCs. We found two such intestine-expressed genes, *acd-3* and *del-5* (Figure 4A, B). Whereas ACD-5 and FLR-1 share the highest homology with mammalian ASIC2, ACD-3 shares the highest homology with mammalian ASIC5 and DEL-5 is more distantly related. mKate::DEL-5 and ACD-3::mKate2 showed a diffuse distribution, with evidence of basolateral, but not apical, localization (Figure 4A, B).

**Figure 4.**
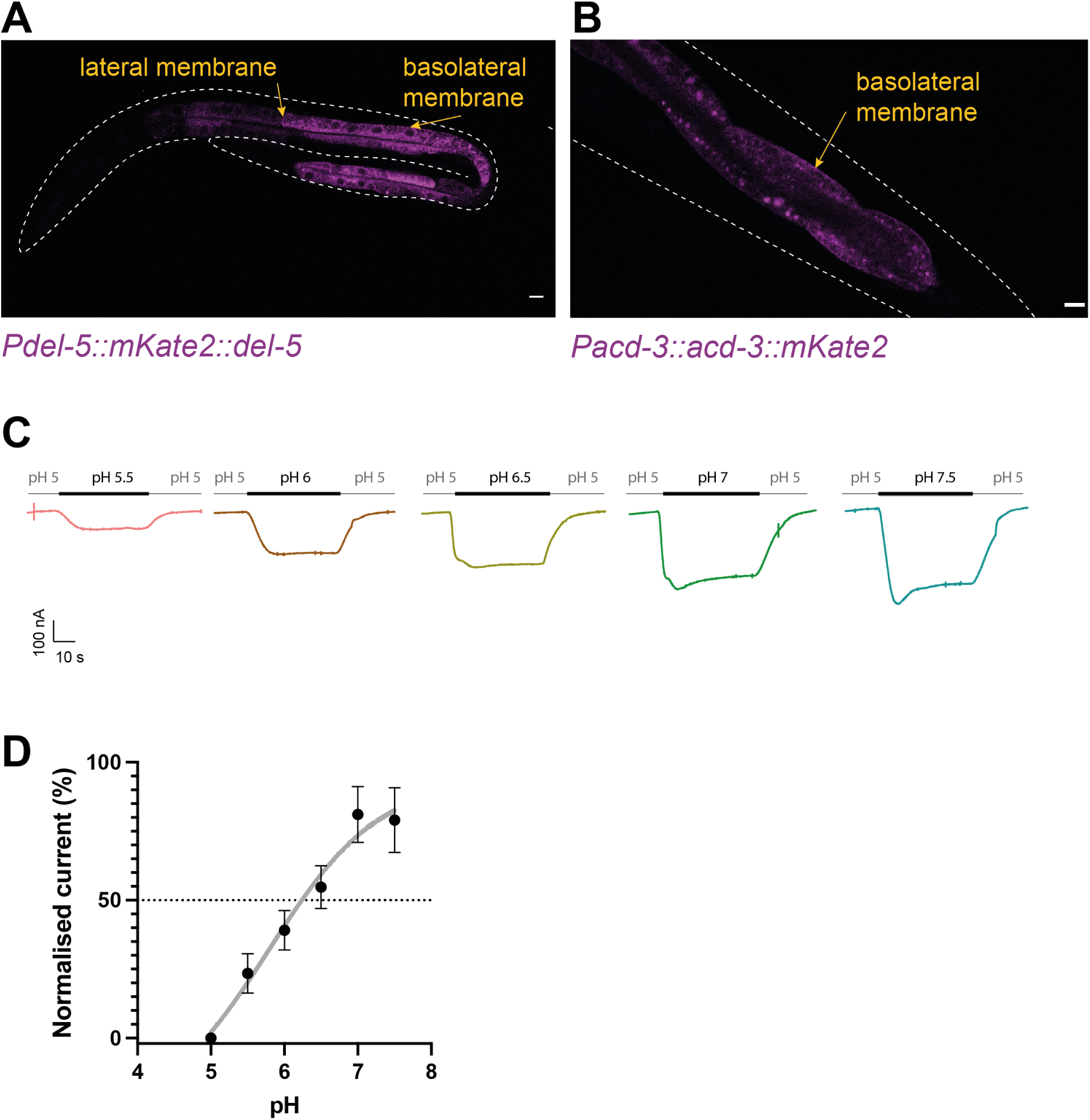
FLR-1 can form pH-sensitive heteromeric channels with ACD-3 and DEL-5. (**A**) ACD-3 is localized at the basolateral intestinal membrane. Localization of mKate2-C-terminally tagged ACD-3 *ljEx1607 (Pacd-3::acd-3 gDNA::mKate2)* (magenta) shows evidence of basolateral membrane localization (yellow arrows). **(B**) DEL-5 is localized at the basolateral intestinal membrane. Localization of mKate2-N-terminally tagged DEL-5 *ljEx1621 (Pdel-5::Nsig::mKate2::del-5 gDNA)* (magenta) shows evidence of basolateral and lateral membrane localization (yellow arrows). Scale bar:10 μm. (**C**) Representative traces from *Xenopus* oocytes injected with *flr-1*, *acd-3* and *del-5* cRNA (250 ng/μL of each construct) for the pH ranges pH 5.5 – pH 7.5 (grey bar represents perfusion time) from a holding pH of pH 5. Currents were recorded at a holding potential of -60mV and traces are baseline-subtracted and drift-corrected using the Roboocyte^2+^ (Multichannels) software. (**D**) pH response curve covering the ranges pH 5 – pH 7.5 from a holding pH of pH 5. N=10. Mean ± SEM. Currents were recorded at a holding potential of -60mV, normalized to maximal currents and best fitted with the Hill’s equation (Nonlin fit Log(inhibitor) vs normalized response – variable slope) in GraphPad Prism.

The presence of these three DEG/ENaC subunits on the basolateral membrane suggested that they might function as heteromers. We therefore asked whether co-expression of ACD-3, DEL-5 and FLR-1 in *Xenopus* oocytes would result in pH sensitive currents. ACD-3, DEL-5 and FLR-1 expressed alone were insensitive to acidic pH (Figure 4-figure supplement 1A, B) and only the DEL-5 current was amiloride-sensitive, in fact showing slightly enhanced currents (Figure 4-figure supplement 2A-C), a phenomenon reported for some mammalian ASICs (*44, 45*). However, when we co-expressed FLR-1 with either DEL-5 and ACD-3, or both, we observed a response to changes in pH from a holding pH 5 (pH_50_ of 5.8), a profile resembling that previously reported for the TadNaCs, DEG/ENaC channels of the marine placozoa *Trichoplax adhaerens* (*36*) (Figure 4C, D, Figure 4-figure supplement 3A, B). We could not detect any currents in response to pH when expressing DEL-5 together with ACD-3 (Figure 4-figure supplement 3C). Thus, FLR-1, together with ACD-3 and/or DEL-5, appears to form a functional acid-inhibited channel with different properties to ACD-5. These findings support the hypothesis that there is a second intestinal DEG/ENaC, a FLR-1, ACD-3 and DEL-5-containing heteromeric channel localized to the basolateral membrane.

### FLR-1, ACD-3 and DEL-5 function together, upstream of Ca*2+*, to control DMP timing

*flr-1(ut11)* mutants show high variability of DMP cycle length, with many extremely short cycles (less than 30 seconds) (*16, 46*) (*16, 33*). We created *flr-1(syb3521),* using CRISPR/Cas9 (*42*), truncating the N-terminus and predicted TM1 with the remaining gene being out of frame so no protein is generated (Figure 1-figure supplement 2C, D). As previously described for *flr-1(ut11)* and other alleles (*16, 43, 46*), *syb3521* presented with a slow-growing, caloric restriction-like phenotype and were thus difficult to cross. Therefore, we used RNA interference (RNAi) by feeding (*47*) to knock down *flr-1* in order to assess genetic interactions with the other potential subunit genes. We used *acd-3(ok1335)*, a truncation of most of the extracellular loop and TMD2, and *del-5(lj138),* generated using CRISPR/Cas9 to delete the N-terminal fragment, the remainder of the gene being out of frame, so no protein is generated (Figure 1-figure supplement 2E-H).

As expected, *flr-1*(RNAi) animals also exhibited the caloric restriction phenotype. Like *flr-1(ut11)*, both *flr-1*(RNAi) and *flr-1(syb3521)* animals showed disruption of DMP, with a dramatic increase in variability, a mix of extremely short and long cycles, missed EMCs and missed or weak pBoc contractions (Figure 5A-D and Figure 5-figure supplement 1). Interestingly, *acd-3/del-5* double mutants phenocopied the *flr-1* deficient animals, showing similar defects in cycle length, EMC and pBoc defects (Figure 5A-D). *flr-1(RNAi)* did not significantly alter these phenotypes, suggesting that the three genes function together and further supporting the hypothesis that these three subunits form a heteromer. Again, we observed a compensatory effect, where a long cycle was followed by a short one (Figure 5-figure supplement 2). Neither of the *acd-3* or *del-5* single mutants exhibited such phenotypes, suggesting that they act redundantly.

**Figure 5.**
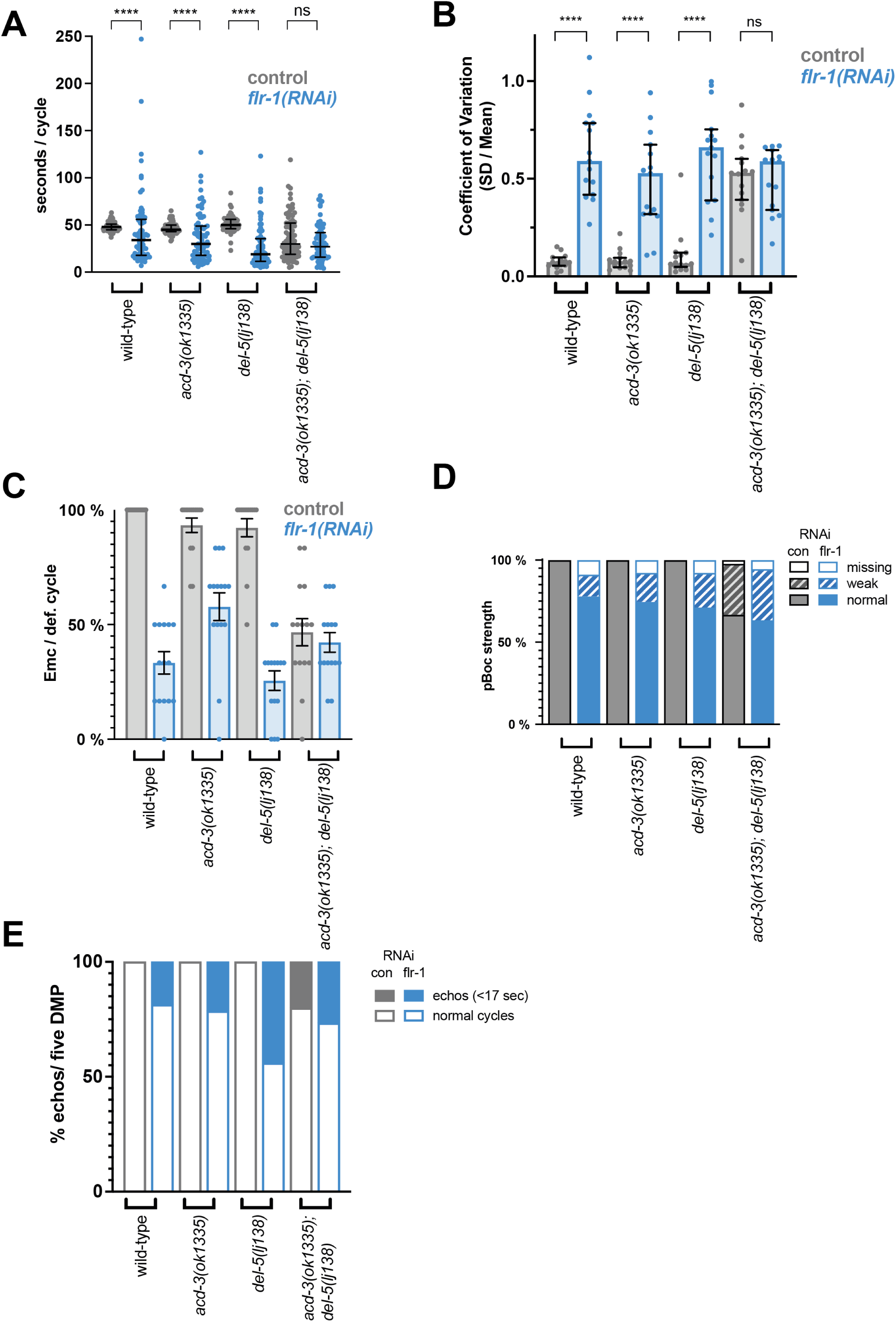
Knock-down of *flr-1* and mutations in *acd-3/del-5* results in arhythmic, missed muscle contractions and echos of the DMP. FLR-1 or ACD-3/DEL-5 depletion alters cycle length and variability. (**A**) Quantification of DMP intervals and (**B**) Coefficient of Variation of animals on *flr-1*(RNAi) (blue) vs control RNAi (grey) assessed by a Mann-Whitney U test (\*\*\*\**p*<0.0001, non-significant (ns)). Error bars represent Median and IQR. (**C-E**) Knock-down of *flr-1* and mutation *acd-3/del-5* result in similar DMP defects. Quantification of EMCs (Mean and SEM) and strength of the pBoc and “echo” DMPs (defined as cycles that last for under 17 seconds (*26, 48*)) of wild-type, *acd-3* or *del-5* single and *acd-3*/*del-5* double mutants on *flr-1(RNAi)* and control (con) RNAi represented as percentages. N=13-15 animals for each genotype x 5 cycles scored for each animal.

Both the *flr-1* deficient animals and the *acd-3/del-5* double mutants frequently exhibited short <17 second cycles, seen for *unc-43(sa200)* CaMK II mutants and described as “echoes,” since they are often incomplete (missing EMC) (*26, 48*) (Figure 5E). The CenGen.org single-cell RNA sequencing database (*49, 50*) indicates that *acd-3* and *del-5* are also expressed in DVB, a motorn neuron involved in initiation of EMCs, so the observed phenotype could also result from neuronal defects. However, previous research has shown that interrupting the Ca^2+^ wave with heparin, an IP_3_ receptor inhibitor, in parts of the intestine eliminates the aBoc and EMC following the pBoc, suggesting that the Ca^2+^ wave is upstream of these steps (*26*). The similarities with both the *itr-1* (highly variable, long cycles), and *unc-43* (echoes) phenotypes suggests that the *acd-3/del-5* and *flr-1* mutations are likely to interfere with the intestinal Ca^2+^ wave.

As Figure 6 shows, *flr-1(syb3521)* completely disrupted the Ca^2+^ oscillations that we normally observe in wild type animals, to the extent that we could not detect obvious peaks. Consistent with the DMP phenotype, we saw some variation, with some animals exhibiting a constant low concentration and others showing more variation; but overall, both the basal and maximal Ca^2+^ concentrations were significantly decreased (Figure 6-figure supplement 1A, B). Taken together, our data indicate that FLR-1 functions in a heteromer along with ACD-3 and/or DEL-5 and that this/these proton gated channels, exposed to the pseudocoelom, play a major role in controlling the intestinal Ca^2+^ transients that are so central to the DMP.

**Figure 6.**
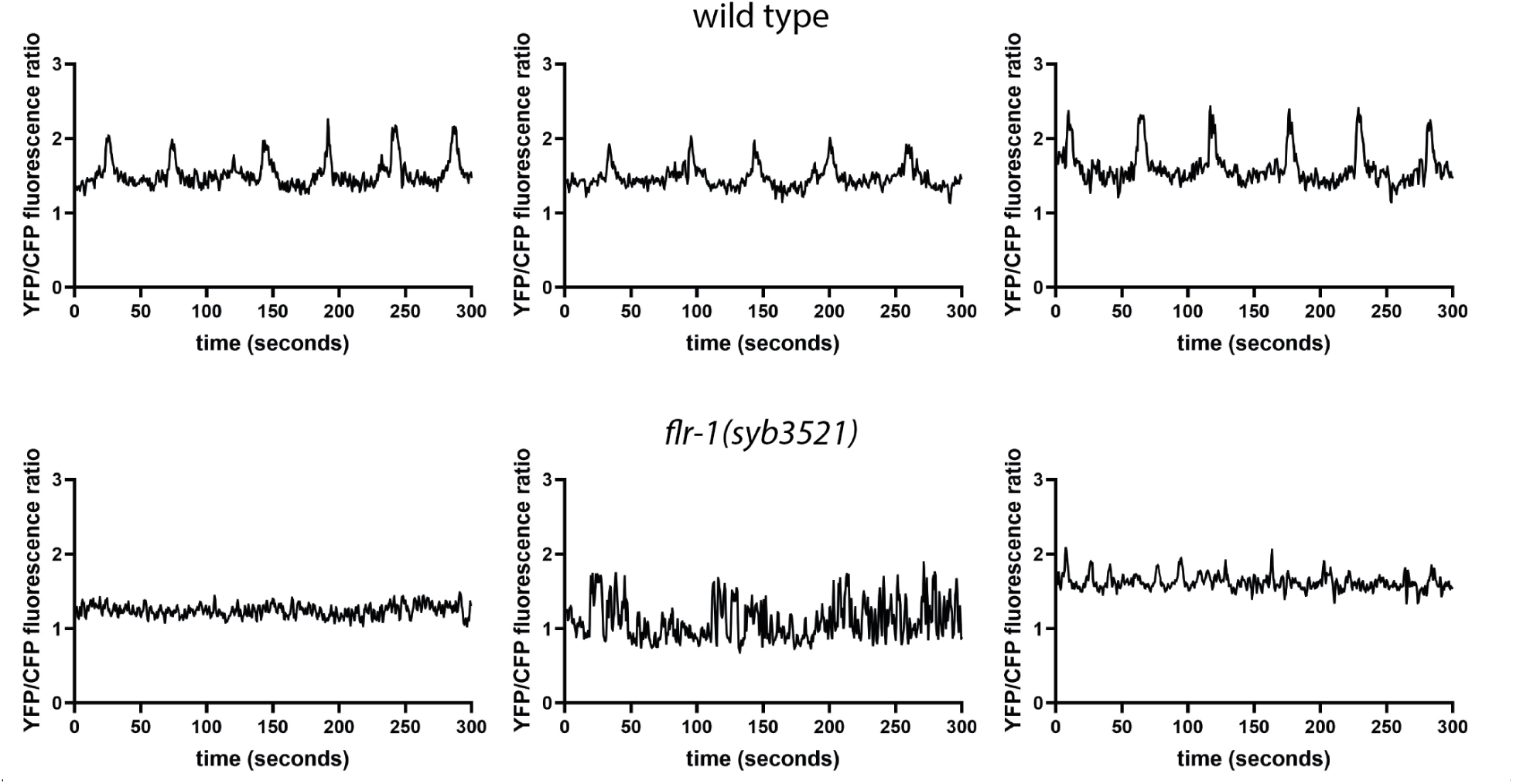
FLR-1 is required for rhythmic intestinal Ca^2+^ transients. Representative example traces of intestinal Ca^2+^ transients, recorded in freely moving animals.

### Dysregulated intestinal proton signaling results in developmental delay and reduced fat storage

Many of the physiological changes associated with dysregulation of intestinal lumen pH reflect starvation, including developmental delay or arrest and reduced growth rate (*23, 27, 31*). These characteristics resemble the phenotypes that have previously been described for *flr-1* mutants (*16*). We noticed that the *acd-3/del-5* mutants also had an appearance suggesting caloric restriction, whereas the single *acd-3* and *del-5* mutants had a superficially normal appearance (Figure 7-figure supplement 1). However, we also wondered whether the disruption of pH homeostasis had consequences for development and metabolism. Therefore, we investigated the physiological consequences of these mutations in more detail. We quantified the time to develop, from laying of the egg to onset of egg laying by the offspring (*51*). Wild-type and the *acd-5(lj122)* mutants showed similar developmental timescales (median 74 and 74 hours), while there was a small but statistically significant delay in developmental timing for the *acd-5(ok2657)*, *acd-3(ok1335)* and *del-5(lj138)* (median 75, 79 and 78 hours, respectively) (Figure 7A). By contrast, the *flr-1(syb3521)* single mutants and the *acd-3/del-5* double mutants showed a severe developmental delay, taking three times as long to reach adulthood (Figure 7B). A small fraction of these animals arrested and died as larvae or before egg-laying. When we quantified the size of day-1 adults, we found that the *acd-3*, *acd-5* and *del-5* single mutants show a wild-type length; by contrast, *flr-1(syb3521)* single mutants and the *acd-3/del-5* double mutants are significantly shorter (Figure 7C). These phenotypes are consistent with a role for FLR-1, ACD-3 and DEL-5 in development and growth.

**Fig. 7.**
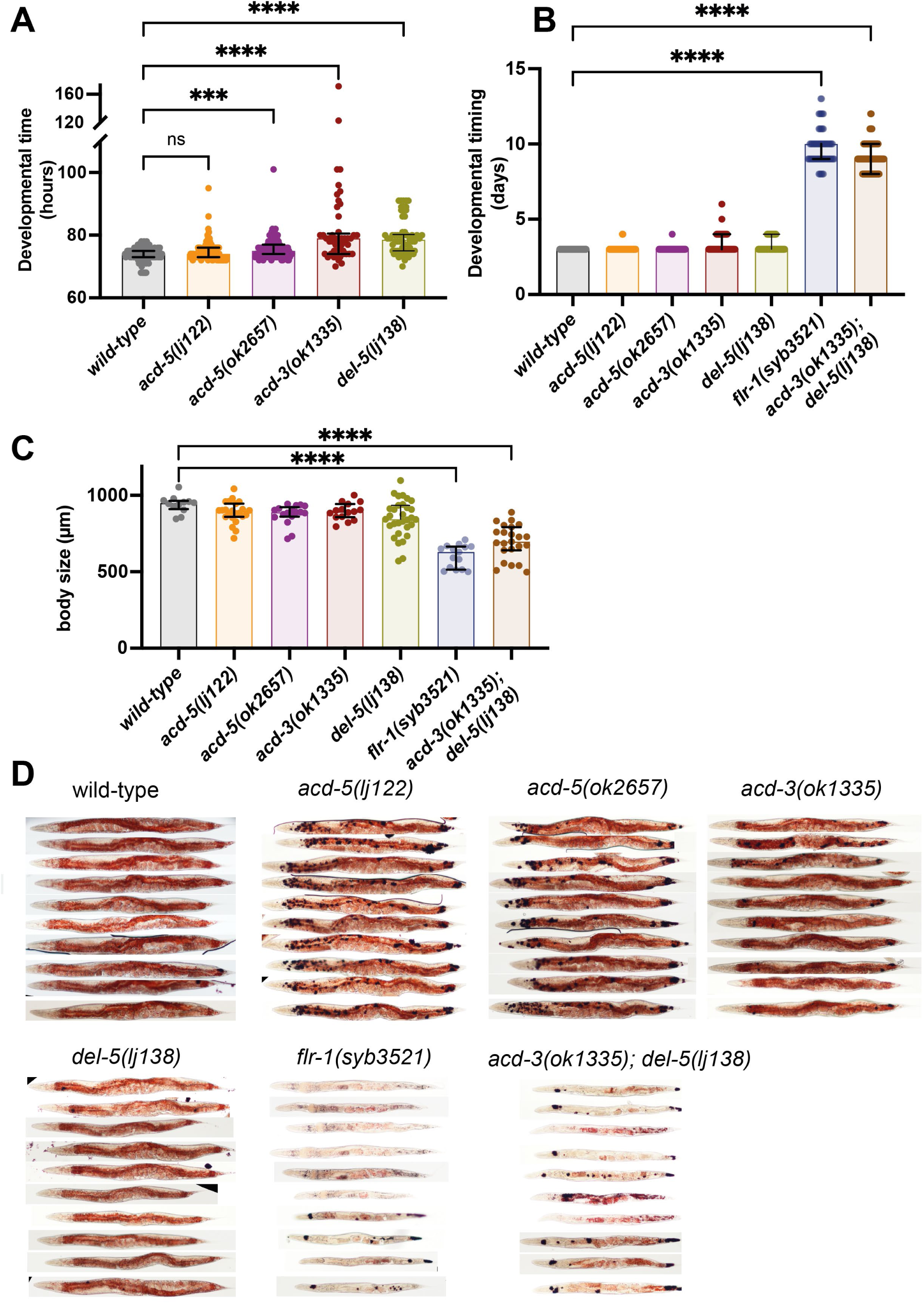
Consequences of disruptions of proton signaling in the intestine. (**A**) *acd-5(ok2657)*, *acd-3* and *del-5* single mutants show a subtle developmental delay. Developmental timing (generation time) of DEG/ENaC single mutants in hours from egg laid to first egg laid as an adult. N= 115, 102, 96, 49, 62, in order on graph. Kruskal Wallis test with Dunn’s multiple comparison post-hoc test, \*\*\**p*=0.0004, \*\*\*\**p*<0.0001. (**B**) *flr-1* single mutants and *acd-3/del-5* double mutants show a severe developmental delay. Developmental timing of DEG/ENaC single, double and triple mutants in days from egg-laid to day of onset of egg-laying. N = as in A for single mutants; 76, 92 for the remainder. (**C**) *flr-1* single mutants and *acd-3/del-5* double mutants are short. Quantification of body size of DEG/ENaC single, and double mutants. Each circle is an individual worm. N=12, 23, 17, 15, 31, 15. Kruskal Wallis test with Dunn’s multiple comparison post-hoc test, \*\*\*\**p*<0.0001. Error bars indicate median and IQR. Each individual point is one animal. (**D**) *flr-1* single mutants and *acd-3/del-5* double mutants show aberrant fat metabolism. Fat storage and distribution of DEG/ENaC single and double mutants assessed by Oil-Red-O staining of 10 representative animals. Scale bar: 50 μm.

Peptide absorption is dependent on proton gradients where nutrients are taken up against the gradient and greatly enhanced at a low extracellular pH (*52–54*). In *C. elegans,* the proton gradient is known to drive nutrient uptake via the OPT-2/PEP-2, a proton-coupled oligopeptide transporter, which is also involved in fat accumulation and *opt-2* mutants show a decrease in intestinal fat deposits (*41*). Previous research revealed a link between DMP defects and aberrant fat metabolism, as well as between dysregulation of intestinal pH and reduced fat storage resulting in nutrient uptake deficiency (*27, 30*). We therefore used Oil-Red O, a fat-soluble dye for staining triglycerides and lipoproteins, to qualitatively observe fat distribution across tissues in the DEG/ENaC mutants (*55*). We found a clear reduction in intestinal fat accumulation, indicated by reduced Oil-Red O staining in *flr-1* single and *acd-3/del-5* double mutants, demonstrating aberrant fat metabolism (Figure 7D). This is consistent with the reduced growth rate and decreased body size and underlines the importance of proper regulation of the DMP, by the FLR-1/ACD-3/DEL-5 channel.

## Discussion

Using a combination of electrophysiology, behavioral analysis, genetic manipulations, and pH and Ca^2+^ indicator imaging, we have obtained evidence for two acid-sensitive DEG/ENaC channels with distinct functions (Figure 8). One includes ACD-5, which localizes to the luminal membrane and is the main proton sensing subunit of the channel. This proton sensor is essential for maintenance and establishment of acidic luminal pH and yet only subtly influences the DMP, and the Ca^2+^ oscillator. The second, composed of FLR-1, ACD-3 and/or DEL-5, localizes to the basolateral membrane and is essential for timing and execution of DMP muscle contractions, with consequences for fat metabolism and growth.

**Figure 8.**
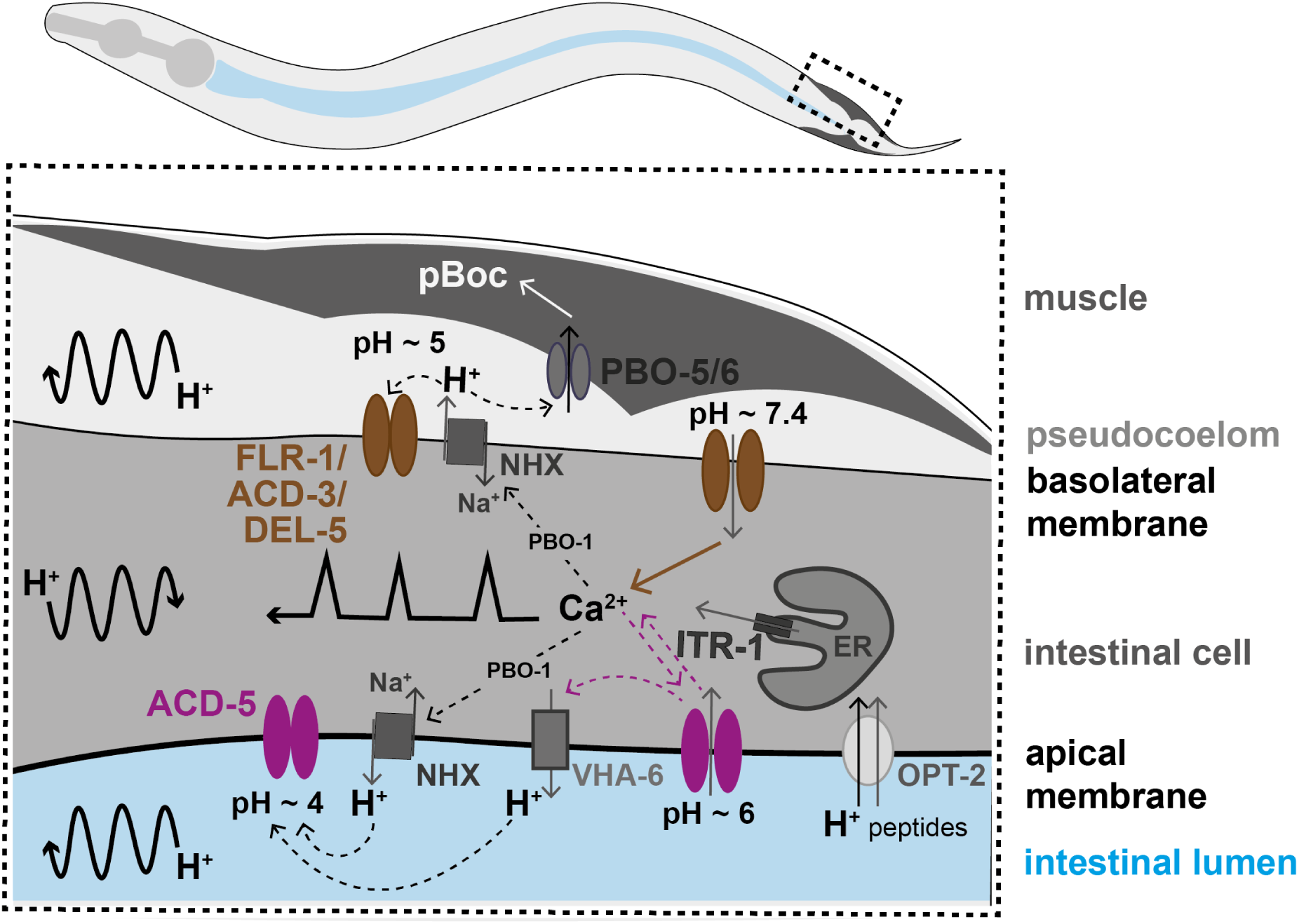
Working model of the two acid-sensing DEG/ENaCs in the *C. elegans* intestinal cells. Ca^2+^ release from the endoplasmic reticulum (ER), via the IP_3_ receptor, ITR-1, is responsible for the Ca^2+^ oscillations transitioning from the posterior to the anterior part of the intestine (represented by the black line with sharp peaks). Ca^2+^ is also responsible for proper localisation of Na^+^/H^+^ exchangers (NHX) via the CHP1 homolog PBO-1. H^+^ oscillations occur in the intestinal cell, transitioning posteriorly, and pseudocoelom and intestinal lumen, transitioning anteriorly (represented by the curved black arrow lines). The FLR-1/ACD-3/DEL-5 containing channel localizes to the basolateral intestinal membrane, it is likely open for much of the cycle, directly affecting Ca^2+^ oscillations and ion homeostasis. The Ca^2+^ transient triggers efflux of H^+^ via NHXs (including PBO-4), decreasing pseudocoelomic pH and closing FLR-1/ACD-3/DEL-5, and Ca^2+^ concentration returns to baseline. ACD-5 is localized at the apical membrane. Proton pumps such as NHXs and VHA-6 maintain the acidity in the lumen during most of the DMP (pH∼4, ACD-5 is closed). Once a cycle, H^+^ influx, through the H^+^/dipeptide symporter OPT-2 and other unknown channels raises the luminal pH to about pH∼6 (ACD-5 opens), the resulting cation-influx reactivates H^+^ efflux (via VHA-6?). ACD-5 has limited influence on the Ca^2+^ oscillations, masked by the dominant role of IP_3_ signalling.

ACD-5 and FLR-1-containing channels exhibit very different pH dependence and kinetics; ACD-5 is fully open at pH 6 and closed at pH 4, reflecting the range of pH oscillations in the lumen (*27*), whereas FLR-1-containing heteromers form a proton-inhibited cation leak channel, similar to ACD-1, TadNaC6 and vertebrate ASIC5 (*34, 36, 56*). This presumably reflects their different physiological contexts, just as the pH dependence of human ENaCs reflect the pH range in epithelia where they are expressed (*57*). Neither ACD-5 nor FLR-1 currents desensitize; the channel remains open for the duration of the pH stimulus (although we see some evidence of partial desensitization of FLR-1 at certain pHs). This contrasts with the fast desensitization of mammalian ASICs and is more analogous to the slower kinetics of ENaCs. It coincides well with our evidence that ACD-5 is actually required for establishment or maintenance of acidic luminal pH; i.e. its activation ensures a return to the condition (pH ∼ 4) at which it is closed. By contrast, the FLR-1/ACD-3/DEL-5 heteromeric channel on the basolateral membrane is predicted to be constitutively open as the extracellular resting pH is around 7.35 (*27*).

All four intestinal DEG/ENaC subunits influence the DMP, but the degree of influence differs dramatically, providing us with mechanistic information about the relationships between extracellular and intracellular events. Research to date indicates that intestinal IP_3_ dependent Ca^2+^ signaling act as the master oscillator and is responsible for cycle length and timing. Disrupting IP_3_ signaling results in a dramatic lengthening and disruption of rhythmicity of the DMP, and a direct relationship is seen between timing of Ca^2+^ fluctuations and the DMP (*19-21, 26, 33*). Our *acd-5* mutant data supports this hypothesis; the underlying Ca^2+^ fluctuations, and DMP rhythmicity, were only subtly affected, despite dramatic disruption of luminal pH, underlining the dominance of the Ca^2+^ oscillator. Where detectable, the MAT interval length was also not affected, suggesting that ACD-5 is not responsible for the timing of the proton wave, but acts downstream. This is supported by the phenotype of animals deficient in the V-ATPase, VHA-6; lumen pH remained more neutral, but robust Ca^2+^ oscillations were maintained, with only a small increase in cycle length (similar to *acd-5(ok2657)*). This suggests a functional link between the two; a pH∼6-activated cation channel and a presumed H^+^ pump whose effect would be to acidify the lumen. Indeed, such a coupling has been proposed in Na^+^ uptake in the Zebrafish larva (*58*). We could speculate that the dominant mutation could result in interference with VHA-6 function, or alternatively it could be interfering with localization or function of the other DEG/ENaC subunits (as we have shown that it can do for ACD-5 itself, when expressed in oocytes). Altering the subunit composition of the FLR-1-containing channels, rather than disrupting it completely, could result in such a subtle extension. Whatever the mechanism, it is clear that the Ca^2+^ oscillator is largely unperturbed. We do nevertheless see some influence; in particular, when the Ca^2+^ oscillator is disrupted, in *itr-1* deficient animals, the very long cycles observed are dependent on *acd-5*, revealing an otherwise constraining role. The Ca^2+^ oscillator is thus upstream, and/or largely independent, of lumen pH, and is the dominant force driving DMP rhythmicity.

In stark contrast to *acd-5* mutants, *flr-1* deficient animals exhibit a dramatic dysregulation of the DMP, with a combination of very long and very short cycles, underlining their importance for rhythmicity (as previous research showed (*16, 33, 43, 46*)), which is phenocopied by *acd-3/del-5* double mutants. The lack of a clear phenotype for *acd-3* and *del-5* single mutants suggests that they act redundantly, in a heteromer, along with FLR-1. This correlates well with our observation that FLR-1 pH-sensitive currents in *Xenopus* oocytes were dependent on the presence of one or other of these subunits, and that *acd-3/del-5* double mutant phenocopy other aspects of *flr-1* loss of function: EMC and pBoC defects, developmental timing, growth, and fat accumulation. The severity of disruption to the DMP is reminiscent of that seen when Ca^2+^ signaling is disrupted (*19–21, 33*). Indeed, we found that in *flr-1(syb3521)* animals, the intestinal Ca^2+^ transients, measured in the posterior cells, where the Ca^2+^ wave initiates, were completely disrupted. This indicates that the FLR-1/ACD-3/DEL-5 channel(s) is upstream of the Ca^2+^ oscillator, perhaps forming part of the “master oscillator”, linking extracellular proton concentration to the initiation of the intracellular Ca^2+^ wave that controls periodicity and execution of the DMP (see Figure 8).

Interestingly, in contrast to *itr-1*, the *flr-1* and *acd-3/del-5* deficient animals also exhibit very short cycles, and “echos” of the DMP as well as missed or weak EMC or pBoc muscle contractions which has previously been described for the *unc-43* CaMK II mutant (*26*) which again links the phenotypes of these mutants to disturbance in intestinal Ca^2+^ signaling. These shorter cycles indicate a role for CaMK II and the FLR-1 channel in retarding a shorter timescale oscillator. Cycle variability, the other effect of *flr-1* or *acd-3/del-5* mutations, presumably represents a loss of synchronicity between semi-independent oscillators, each of which has an innate homeostatic rhythm – but is checked by a delay in reaching permissive conditions provided by another. A channel that is open for most of the cycle is consistent with a role in maintaining cellular excitability, and thereby underpinning intracellular Ca^2+^ signaling; closing briefly following proton release into the pseudocoelom would serve to terminate the signal. This model fits well with the temporal relationship observed (*27*), where Ca^2+^ concentration precisely follows the decline in pseudocoelom pH.

Notwithstanding the challenges associated with imaging freely moving animals, one important barrier to dissecting the relationships between events in the intestinal cells and their extracellular environments is that some components are expressed on both the apical and basolateral membranes. PBO-1, for example, localizes to both membranes, where it has distinct roles (*59*). Our data for ACD-3 and DEL-5 localization to the basolateral membrane supports previous evidence for FLR-1 (*43*). We did not detect apical localization for any of ACD-3 or DEL-5 subunits. However, Take-uchi et al. (1998) reported apical localization (*16*), so we cannot exclude the possibility of a dual role at the two membranes. Basolateral localization is consistent with the weak and missing pBoc resulting from disturbance in Ca^2+^ and proton homeostasis.

We have shown that two acid-sensing DEG/ENaC channels with distinct properties play very different roles in the interplay between extracellular pH and intracellular Ca^2+^. ACD-5 is required to maintain acidic lumen pH and has a subtle impact on the Ca^2+^ oscillations that control DMP timing. In contrast, a FLR-1, ACD-3 and/or DEL-5-containing channel is the essential link between pseudocoelomic pH and Ca^2+^, and thus timing and execution of the DMP. DEG/ENaCs are conserved across the animal kingdom, and the ASICs and ENaC members are relevant to a broad range of medical conditions, and thus of significant therapeutic importance. Our data provides an insight into the relationship between the acid-sensing properties of these channels and their physiological role in epithelia. However, the family contains a diversity of members with distinct properties, and vastly expanded numbers in some species. Better understanding the relevance of channel properties to function will help us to better understand their role, and the role of acid sensing, in other cellular contexts.

## Materials and Methods

### KEY RESOURCES TABLE

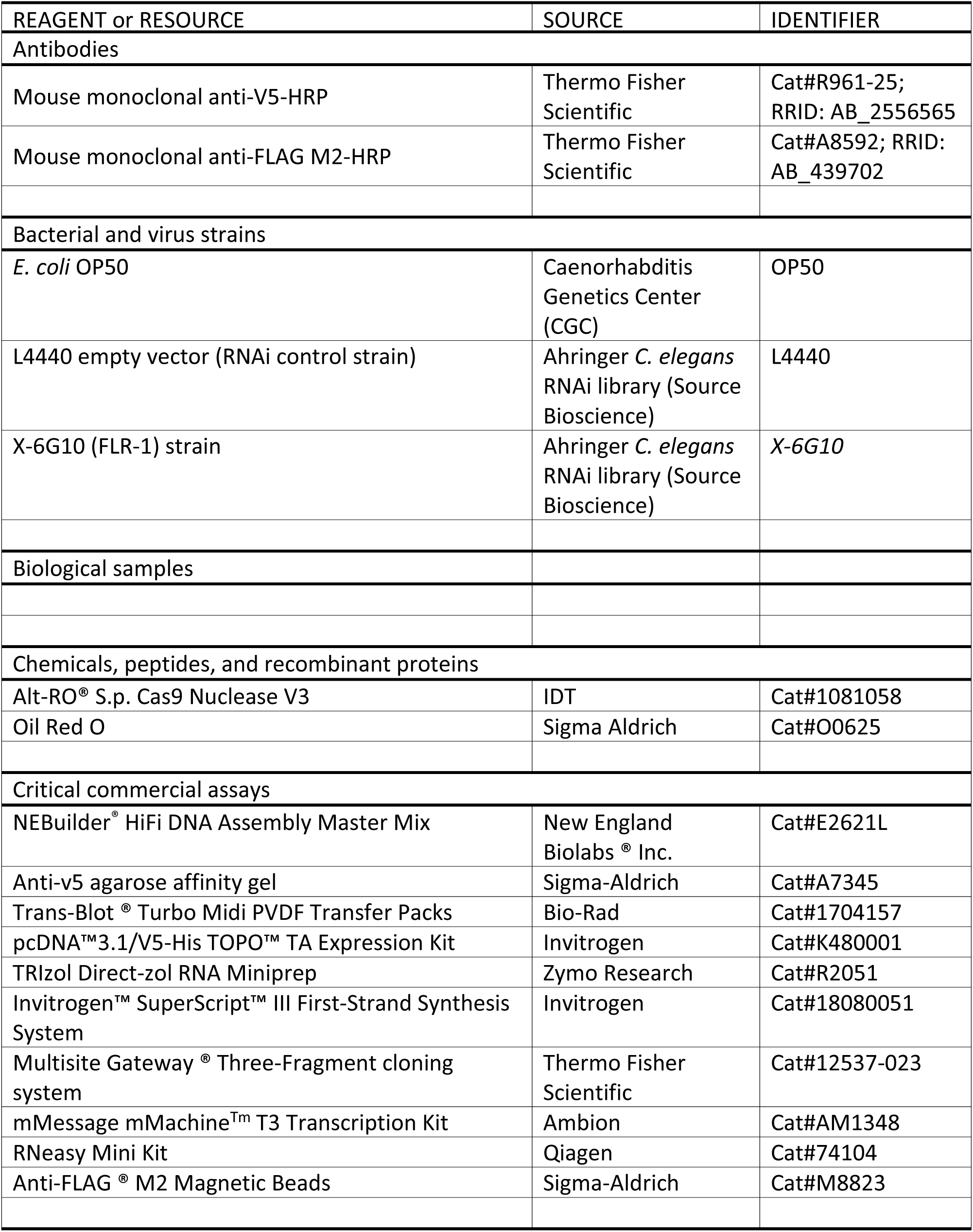
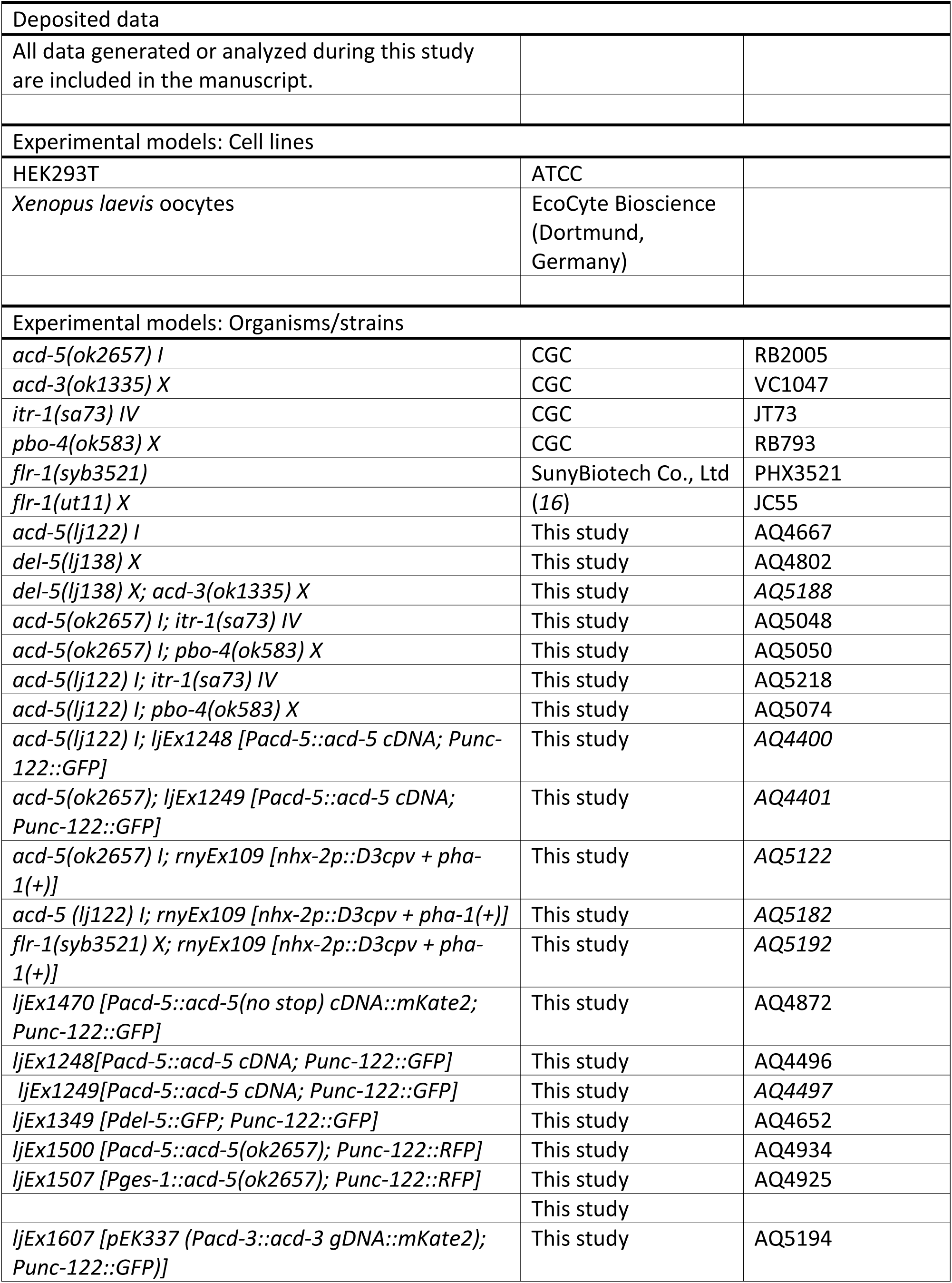
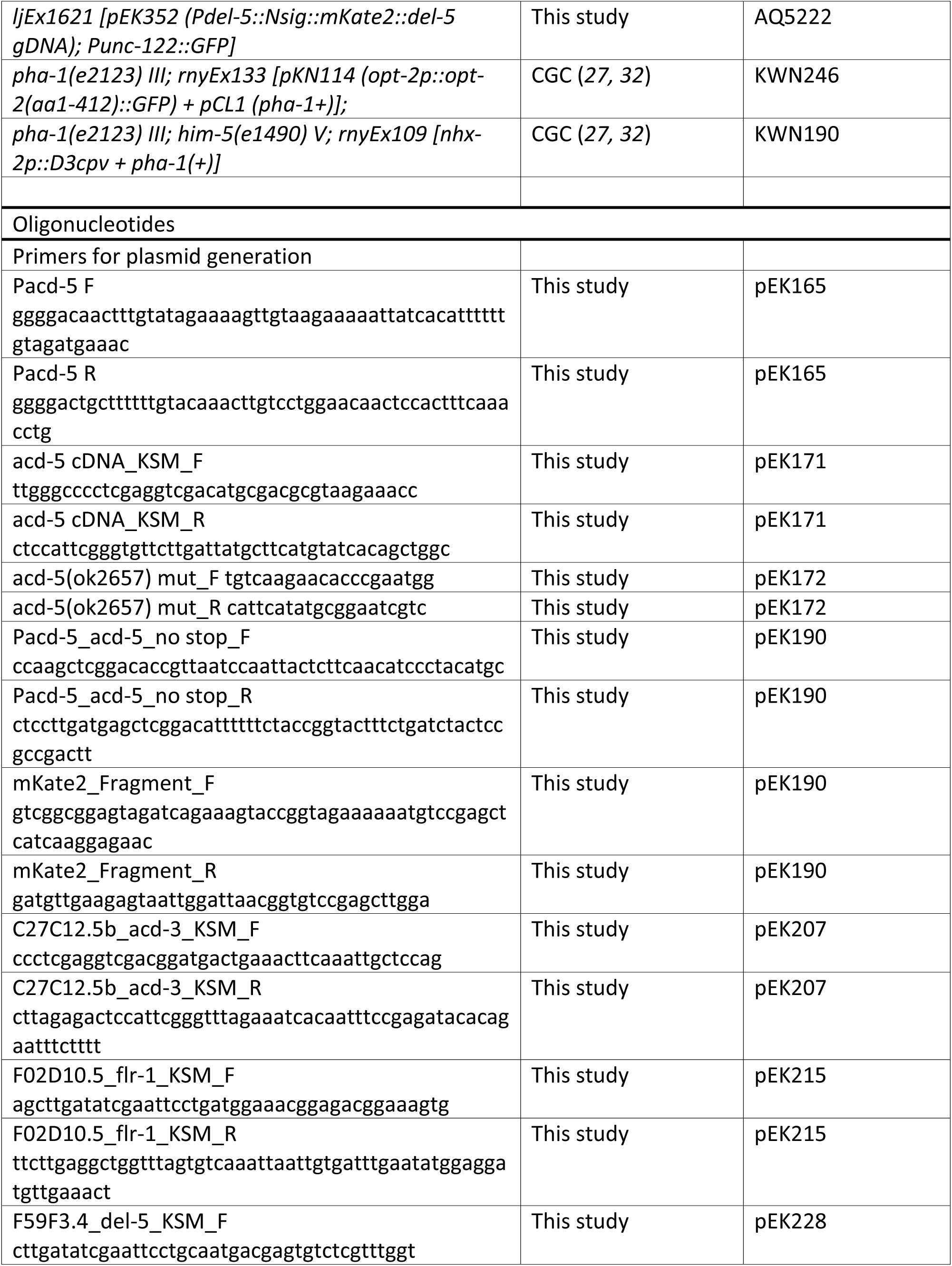
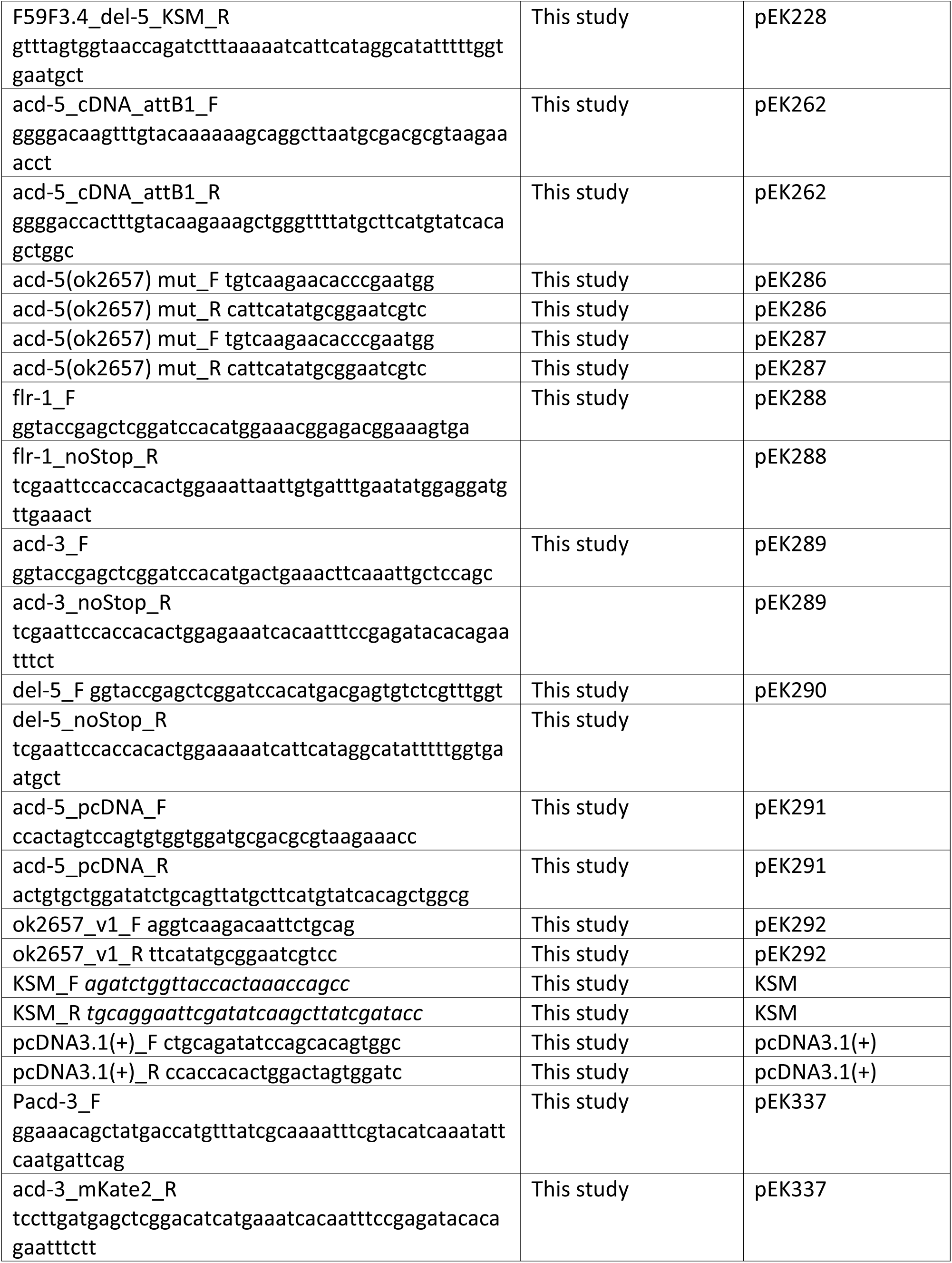
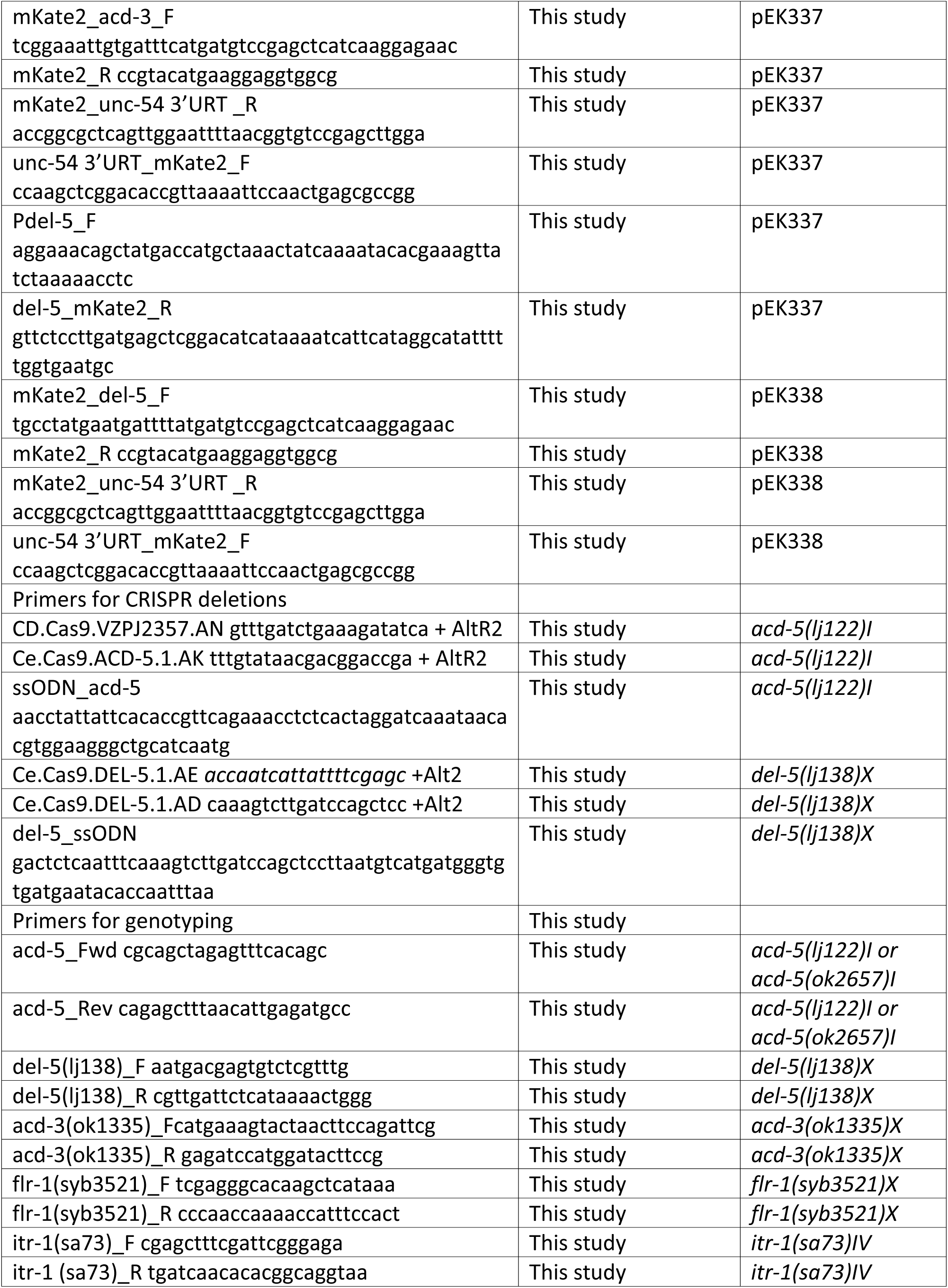
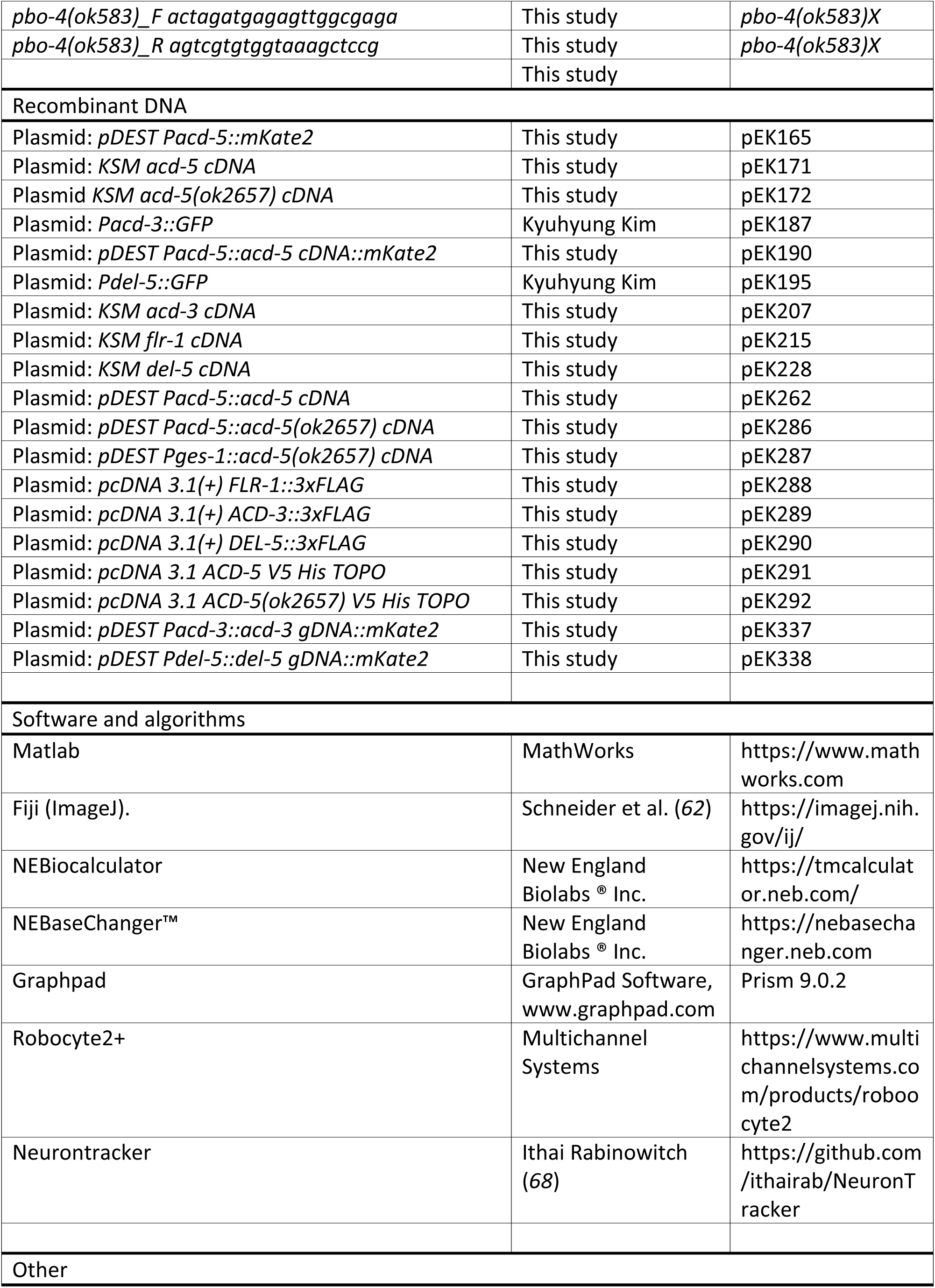
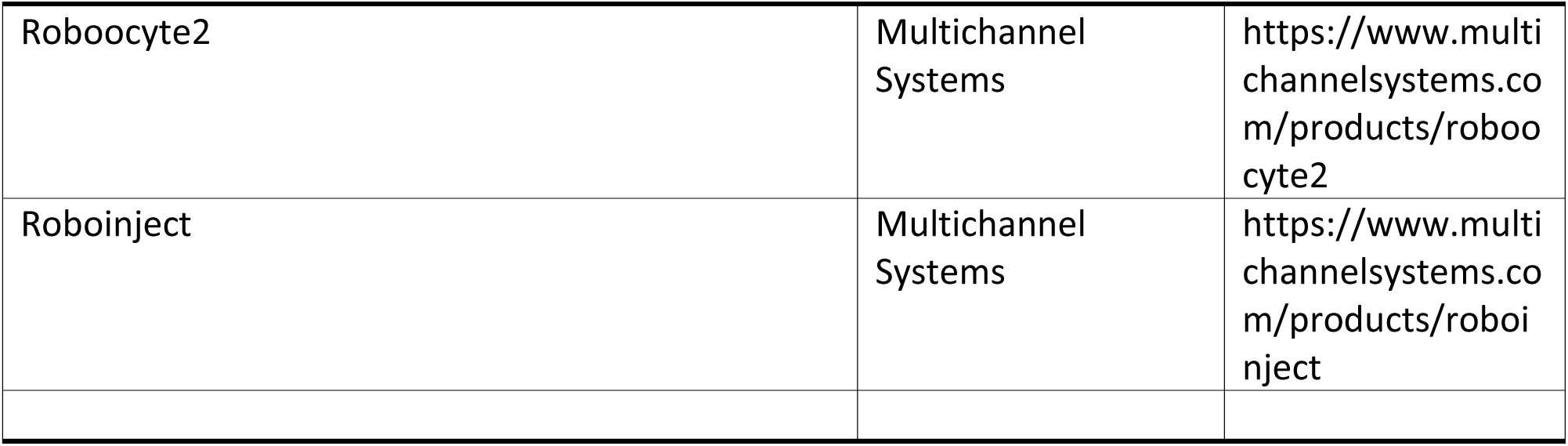

#### *C. elegans* growth and maintenance

Standard techniques were used for *C. elegans* strain maintenance and genetic manipulations (*60*). All experiments were performed on hermaphrodite worms grown on *E.coli* OP50 at room temperature (22°C) and all animals were cultivated at 22°C unless otherwise stated. Strains with the temperature sensitive allele *itr-1(sa73)* were cultivated at the permissive temperature of 15°C to avoid sterility. Mutations were 6x outcrossed with the Bristol N2 wild-type, and transgenic strains were generated by microinjection of plasmid DNA (*61*). In order to generate the CRISPR mutations *acd-5(lj122)* or *del-5(lj138)*, the protocol established by the Mello lab was used (*42*). Guide RNAs (crRNA) and homology arms were ordered from IDT (Leuven Belgium) or Sigma-Aldrich (Merck Life Science UK Limited, Dorset, UK). For subcellular localization of ACD-5 to the apical membrane localization the strain KWN246 *pha-1(e2123) III; rnyEx133 [pKN114 (opt-2p::opt-2(aa1-412)::GFP) + pCL1 (pha-1+)]* was used. The following transgene was used for the free-moving *in vivo* Ca^2+^ imaging experiments: *rnyEx109 [nhx-2p::D3cpv + pha-1(+)]*constructed by Keith Nehrke (*27, 32*). The following strains were provided by the CAENORHABDITIS GENETICS CENTER CGC, which is funded by NIH Office of Research Infrastructure Programs (P40 OD010440): RB2005 *acd-5(ok2657)I;* VC1047 *acd-3(ok1335) X;* JT73 *itr-1(sa73) IV;* RB793 *pbo-4(ok583) X;* KWN246 *pha-1(e2123) III; rnyEx133 [pKN114 (opt-2p::opt-2(aa1-412)::GFP) + pCL1 (pha-1+)];* KWN190 *pha-1(e2123) III; him-5(e1490) V; rnyEx109 [nhx-2p::D3cpv + pha-1(+)].* The following strain was created by SunyBiotech Co., Ltd (Fu Jian Province, China): PHX3521 *flr-1(syb3521)* (1416 bp deletion).

#### Molecular Biology

The transcriptional reporters *Pdel-5::GFP* and *Pacd-3::GFP* plasmids were a kind gift from Professor Kyuhyung Kim’s lab (Daegu Gyeongbuk Institute of Science & Technology (DGIST), Korea) containing regulatory sequences 888 bp upstream of the *del-5* gene and 3085 bp upstream of the *acd-3* gene, respectively. For all other plasmids including the tagged proteins and sub-cloning of cDNA into KSM vector for *Xenopus* oocyte expression, the NEBuilder HiFi DNA Assembly Reaction Protocol was used to assemble the vector and inserts using NEBuilder^®^ HiFi DNA Assembly Master Mix (Catalogue # E2621L) and a vector:insert ratio 1:2 (0.5pmol vector and the corresponding amount of insert using NEBiocalculator) see above. *C. elegans* cDNA was obtained from growing N2 wild-type animals on fifteen 6cm NGM plates until the food was diminished, and subsequently extracted and purified using the TRIzol Direct-zol RNA Miniprep (Catalogue #R2051, Zymo Research). cDNA was generated using the Invitrogen™ SuperScript™ III First-Strand Synthesis System (Catalogue # 18080051). Primers were designed using SnapGene 5.0.4. (Hifi-Cloning of two fragments) based on the cDNA gene sequence found on wormbase.org, and ordered from Integrated DNA Technologies Inc (IDT) (Leuven Belgium) or Sigma-Aldrich (Merck Life Science UK Limited, Dorset, UK). The cDNA inserts were sub-cloned into the KSM vector under the control of the T7 promoter containing 3’ and 5’ untranslated regions (UTRs) of the *Xenopus* beta-globin gene and a poly(A) tail. The forward primer *agatctggttaccactaaaccagcc* and reverse primer *tgcaggaattcgatatcaagcttatcgatacc* were used to amplify the KSM vector. NEB T_m_ Calculator was used to determine annealing temperatures. For generation of mutations and deletions, the NEBaseChanger™ tool to generate primer sequences and an annealing temperature. The *acd-5(ok2657)* mutant cDNA was based on the *acd-5(ok2657)* mutant allele generated by *C. elegans* Gene Knockout Consortium (CGC).

#### Confocal microscopy

Worms were mounted on 3% agar pads (in M9 (3 g KH2PO4, 6 g Na2HPO4, 5 g NaCl, 1M MgSO4)) in a 3 μL drop of M9 containing 25 mM sodium azide (NaN_3_, Sigma-Aldrich). Images were acquired using a Leica TCS SP8 STED 3X confocal microscope at 63x, 40x or 20x resolution and Z stacks and intensity profiles were generated using Fiji (ImageJ) (*62*).

#### Two-Electrode Voltage Clamp in *Xenopus* oocytes

Linearized plasmid DNA was used as the template for *in vitro* RNA synthesis from the T7 promoter using the mMessage mMachine T3 Transcription Kit (Ambion # AM1348), producing 5’ Capped RNA. The reaction was incubating at 37°C for 2h, and the resulting RNA was purified by GeneJET RNA Cleanup and Concentration Micro Kit (Thermo Scientific # K0841) and eluted in 15μL RNase free water. *Xenopus laevis* oocytes of at least 1mm in size were obtained from EcoCyte Bioscience (Dortmund, Germany). They were de-folliculated by collagenase treatment and maintained in standard 1xND96 solution (96mM NaCl, 2mM MgCl_2_, 5mM HEPES, 2mM KCl, 1.8mM CaCl_2_, pH7.4. Oocytes were injected with 25μL of cRNA solution at a total concentration of approximately 500 ng/μL using the Roboinject (MultiChannel Systems). Oocytes were kept at 16°C in 1xND96 prior to TEVC. TEVC was performed 1-2 days post injection at room temperature using the Roboocyte2 (MultiChannel Systems). *Xenopus* oocytes were clamped at −60 mV, using ready-to-use Measuring Heads from MultiChannel Systems filled with 1.0 M KCl and 1.5 M K-acetate. All channels were tested using the Roboocyte2 (MultiChannel Systems). For all current-voltage steps (I-V) experiments, measurements were obtained in each solution once a steady-state current was achieved and the background leak current was subtracted.

As millimolar concentrations of Ca^2+^ and other divalent ions except Mg^2+^ can block ASIC currents (*38*), Ca^2+^-free buffers were used for substitution experiments of monovalent cations adapted from a previous protocol (*37*): 96mM XCl, 1mM MgCl^2^, 5mM HEPES, pH adjusted to 7.4 with XOH, where X was Na^+^, K^+^ or Li^+^, respectively. For testing ion permeability for Ca^2+^, a previous protocol was used (*34*) replacing Na^+^ with equimolar Ca^2+^. If necessary, D-Glucose was used to adjust osmolarity. The osmolarity was checked and confirmed to be within the error of 210 mosm (*63*). For testing pH sensitivity, 1x ND96 solutions was used; for solutions with a pH 5 or lower, MES was used instead of HEPES and adjusted with HCl. I-V relationships for ion selectivity were calculated by subtracting the background leak current in the presence of 500 μM amiloride from the current observed in the absence of amiloride in order to get the actual current. Actual current IV curves for each individual oocyte were fitted to a linear regression line and the x intercept was compared between solutions to calculate an average reversal potential (E_rev_). Reversal potential shift (ΔE_rev_) when shifting between pHs or from a NaCl to a KCl, LiCl or CaCl_2_ solution was calculated for each oocyte. In order to test the responses to pH, the channel-expressing *Xenopus* oocytes were perfused with 1x ND96 (using HEPES for buffering pH above 5.5 and MES for pH below 5), pH was adjusted with HCl and ranged from pH 7.4 (neutral pH of the ND96 solution) to pH 4. Background currents measured at pH 7.4 for ACD-5 and pH 5 for FLR-1-containing channels were subtracted from those measured during activation of the channels. For analysis, currents were normalized to maximal currents (I/I_max_) and best fitted with the Hill’s equation (Variable slope).

#### Developmental timing of DEG/ENaC mutants and wild-type animals

The protocol described by (*51*) was used: Briefly, animals were synchronized by allowing adults to lay eggs for one hour on an *E. coli* OP50 seeded 60mm NGM plate. After hatching animals were randomly picked onto individual 20 μL *E. coli* OP50 seeded NGM plates. Time (in hours and days) was counted from laid egg to egg-laying adulthood. The animals were kept at 20°C for the duration of the assay.

#### Assessing and Scoring Defecation Motor Program (DMP)

Defecation was assayed as previously described (*4*). Briefly, animals were synchronized by allowing 10 day-1 hermaphrodites to lay for ∼ 3 hours before picking them off and growing today-1 adult. On the day prior to the assay transgenics were picked to a new plate (where applicable) and blinded. Following 2 minutes acclimation on the microscope, five DMP cycles per animal were observed on a dissecting stereomicroscope at 50X magnification. The time elapsed between two pBocs was measured as one DMP cycle and success of the pBoc and EMC steps was recorded. “echo” DMPs defined as cycles <17 seconds as previously described (*48*). At least 15 animals, assayed across at least 3 days for 5 cycles each and wild-type and mutants were always scored alongside.

RNAi feeding experiments were performed as described (*47, 64, 65*). Standard NGM plates were prepared, with 100 μg/mL carbenicillin and 1mM IPTG added after autoclaving, and were poured 1-3 days before use (*66*). For feeding RNAi experiments the L4440 empty vector was used as a negative control and the X-6G10 (FLR-1) bacterial strain from the Ahringer *C. elegans* RNAi library (Source Bioscience) was used to knock-down the *flr-1* gene. The bacterial strains were freshly grown overnight in LB + 50 µg/mL ampicillin and 24 hours prior to each assay plates were seeded with 100 μL of overnight bacterial culture. Due to the severe developmental delay, animals were picked as L4s and scored as day-1 adults.

#### Imaging and analysis of intestinal lumen pH

KR35 feeding, imaging and image analysis was carried out as previously described (*28, 31*). Briefly, worms were raised to young adults on *E. coli* OP50. Prior to acquisition of videos, the animals were transferred to NGM plates supplemented with 10 μM KR35 and *E. coli* OP50 for 15–30 minutes, and then transferred and imaged on NGM plates without the fluorophore. All animals were treated with equivalent conditions on all days of imaging, including feeding the KR35 dye to animals of each genotype from the same source plate (one condition at a time) to avoid differences in dye concentration, immediately prior to imaging.

Imaging and image analysis were done as previously described (*28, 31*). Videos of free-moving animals fed with KR35 dyes were acquired on a Leica M165FC microscope using a Leica DFC3000G CCD Camera via the Leica Application Suite software (v 4.40) (Leica Microsystems (Switzerland) Limited). Image sequences were acquired at 5x zoom, with 10x gain, and at 10 frames per second. Illumination was via a Leica Kubler Codix source equipped with an Osram HXP 120W lamp. Images were opened in Fiji (ImageJ), converted to 8-bit, scaled from 0–255 to a dynamic range of 10–120, and a rainbow RGB look-up table applied. Movies were converted to AVI using FIJI based on an approximately 30 s clip that corresponded to ∼10–15 seconds before and after a Maximum Anterior Transitions (MAT) (where one occurred). Movies were acquired for ∼2 minutes, to include 1–3 MATs. Movies were opened in ImageJ, and a circular ROI of 25×25 pixels was used to measure the fluorescence of the anterior-most intestine during a MAT. 1-3 measurements were taken per animal, and the data transferred to Excel and GraphPad Prism for graphing and statistical analysis.

#### Ca^2+^ imaging

Ca^2+^ imaging was performed on freely moving animals. Transgenic day-1 adults expressing the genetically encoded calcium indicator Cameleon D3cpv in the intestine were imaged on standard 60mm NGM plates for 5 minutes. Plates had been seeded with 5µL *E. coli* OP50 the day before and grown at room temperature to yield a small lawn, spatially confining the worm’s movement to ease tracking. Animals were picked onto these plates at least 10 minutes prior to recording and were allowed to acclimatize for at least 5 min following mounting of the plate on the imaging microscope, visually confirming that they were defecating, and were manually tracked during video acquisition. Filter/dichroic pairs were as described previously (*67*) Images were recorded at 2 Hz, with 2×2 binning, using an iXon EM camera (Andor Technology), captured using IQ1.9 software (Andor Technology) and analyzed using NeuronTracker, a Matlab (MathWorks) analysis script written by Ithai Rabinowitch (*68*). CFP/YFP fluorescence ratio was smoothed using a 2-frame running average to generate the individual traces shown. Ca^2+^ “peaks” were identified using a 10-frame running average to identify the frame with maximum fluorescence and this information was used to align cycle transients for mean traces, and to calculate the time interval between peaks.

#### Oil-Red-O staining and quantification of worm length

Oil-Red-O staining was performed as previously described (*55*): Day-1 adults were rinsed with 1x PBS, collected in 1.5 mL previously autoclaved Eppendorf tubes (to prevent worms from sticking to the tube), and washed twice with 1x PBS. After the final wash, supernatant was reduced to 120 μL and an equal volume of 2x MRWB (160 mM KCl, 40 mM NaCl, 14 mM Na2EGTA, 1 mM spermidine-HCl, 0.4 mM spermine, 30 mM Na-PIPES pH 7.4, 0.2% β-mercaptoethanol) was added along with 2% paraformaldehyde (PFA). Samples were incubated at room temperature on rocking stage for 1 hour. Worms were then allowed to settle by gravity and washed with 1x PBS to remove PFA. Supernatant was removed and to dehydrate the samples, 60% isopropanol was added and incubated at room temperature for 15 minutes. Oil-Red-O stock solution was prepared beforehand by dissolving the dye in isopropanol to a stock concentration of 0.5 mg/mL. Before use, Oil-Red-O was diluted with water to reach a final concentration of 60%, filtered, and 1 mL of dye added to each sample. Samples were then incubated on a rocking stage at room temperature overnight. Dye was then removed and worms washed with 1x PBS prior to imaging. Animals were mounted on 3% agarose pads and imaged at 10x or 20x magnification using an OLYMPUS BX41 with DIC optics connected to a NIKON Digital Sight. Image straightening was performed using ImageJ.

#### Statistical Analysis

Statistical Analysis was performed in GraphPad Prism version 9.0.2 for macOS, GraphPad Software, San Diego, California USA (www.graphpad.com). Normality (if the data follows a normal/Gaussian distribution) was assessed by a Shapiro-Wilk test. If the data followed a normal distribution, a parametric test was deployed, if not, the non-parametric equivalent was chosen. Appropriate post-hoc tests were always used for multiple comparisons (Bonferroni correction). The statistical tests used are indicated in each figure description.

## Acknowledgements

We are very grateful to Yee Lian Chew, Soudabeh Imanikia, Rebecca Taylor, and members of the Schafer, Taylor and de Bono (LMB), Beets and Timmerman (KU Leuven) and Pless (University of Copenhagen) labs, as well as Ewan St. John Smith (University of Cambridge) for helpful discussions and advice. We are grateful to the LMB Science and Support Facilities, in particular Ben Sutcliffe, Jonathan Howe, and Nick Barry from the Light Microscopy Team for their advice and help with microscopy, and Sue Hubbard, Mark Cussens, and Martyn Howard and the Team from Media and Glass for preparing solutions and NGM plates. We thank Kyuhyung Kim’s lab (Daegu Gyeongbuk Institute of Science and Technology) for providing us with their *C. elegans* DEG/ENaC transcriptional reporter plasmids and sharing their unpublished expression data with us. Some strains were provided by the Caenorhabditis Genetics Center, which is funded by NIH Office of Research Infrastructure Programs (P40 OD010440), and some strains were generated by SunyBiotech (Fuzhou, China). Neurontracker was written by Ithai Rabinowitch. This work was funded by the Medical Research Council (MC-A023-5PB91), Wellcome Trust (WT103784MA) and the KU Center for Chemical Biology of Infectious Diseases (P20GM113117).

## Author contributions

EK, WRS and DSW conceived the project. EK performed and analyzed all experiments, except: DSW performed and analyzed the Ca^2+^ imaging experiments; BDA performed the pH imaging experiments, which EK analyzed. IH and Y-QT provided significant technical guidance for electrophysiology. EK and DSW wrote the manuscript, with contributions from BDA, WRS, IH and Y-QT. All authors contributed to the conception and design of experiments and discussion of results, and approved the final manuscript.

---

**Figure 1-figure supplement 1.**
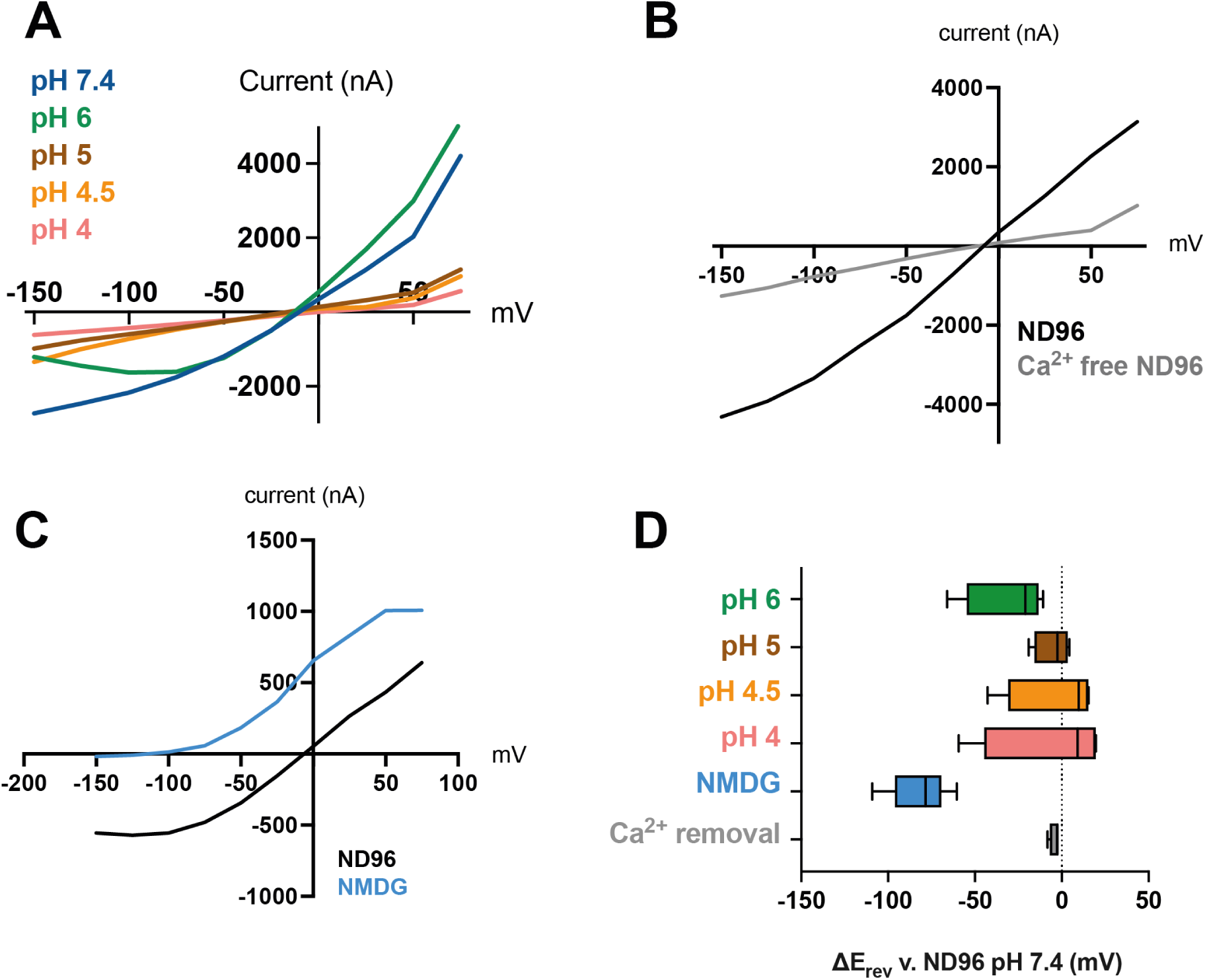
Further characterization of ACD-5 homomeric channel properties. (**A**) Representative current-voltage relationships perfusing different pH solution over the oocyte. (**B**) Representative current-voltage relationships after removal of Ca^2+^ from the solution. (**C**) Representative current-voltage relationships in a basal solution (ND96, pH 7.4) and after removal of Na^+^ from the solution (NMDG, pH 7.4). (**D**) Summary of reversal potential E_rev_ of data presented in A-C. dashed line represents the E_rev_ for the Na^+^ solution. N (top to bottom) = 4, 4, 4, 4, 8, 4. Data is presented as boxplots with Median and IQD (inter-quartile distance, as calculated by the Tukey method).

**Figure 1-figure supplement 2.**
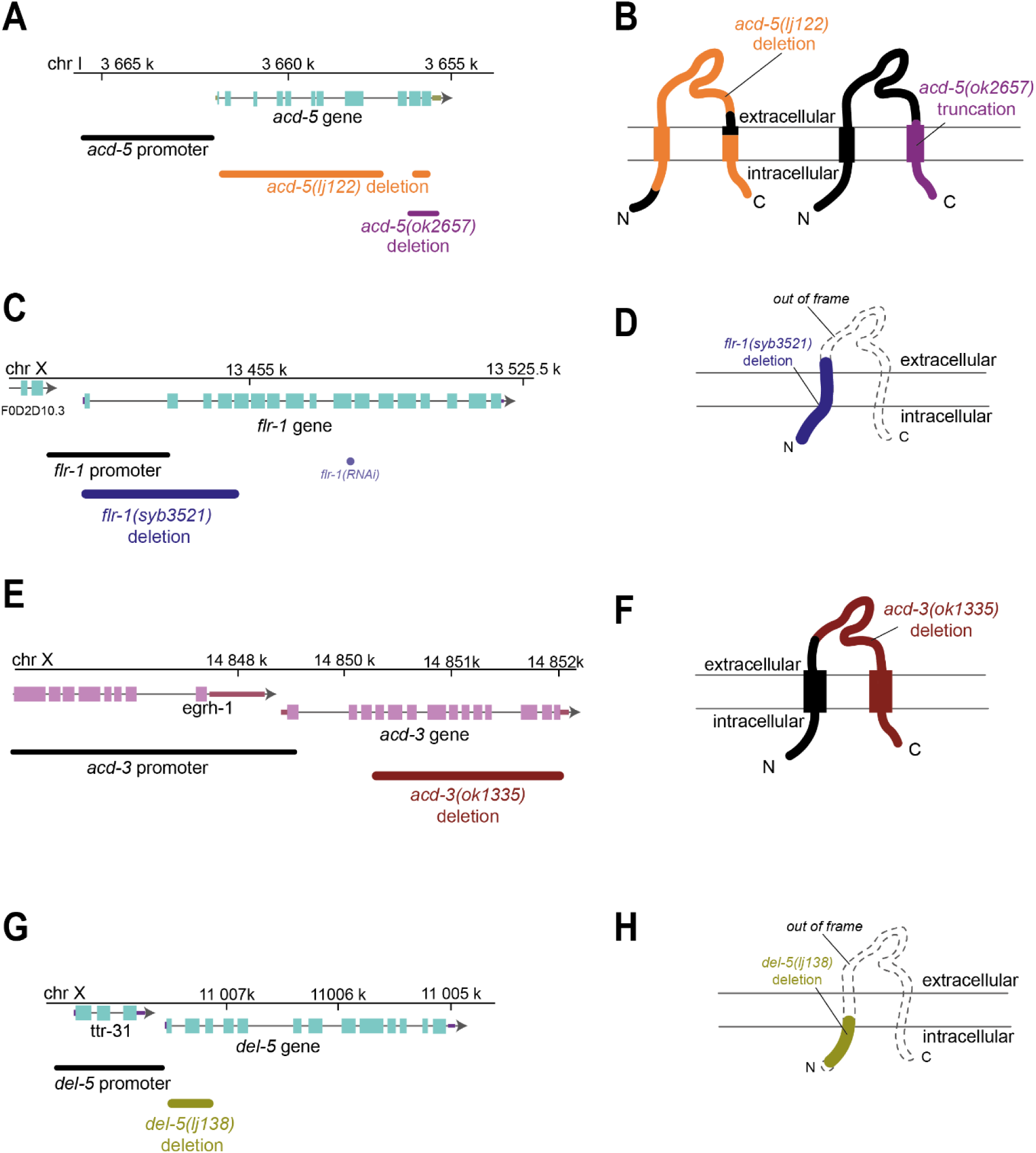
Genomic regions and schematic of predicted protein structures of the intestinal DEG/ENaCs, showing mutations used in this study. (**A, C, E, G**) genomic region for the genes indicated. Boxes indicate exons, lines indicate introns, chr indicates chromosome. Endogenous promoter region used, mutations and RNAi used in this study are shown. (**B, D, F, H**) Schematic of predicted protein structure with two transmembrane domains, and cytosolic N- and C-terminus and an extracellular loop, typical of DEG/ENaC subunits. Mutations used in the study are indicated for each gene and protein. Black indicates intact; colored indicates deleted; dashed line indicates out of frame.

**Figure 1-figure supplement 3.**
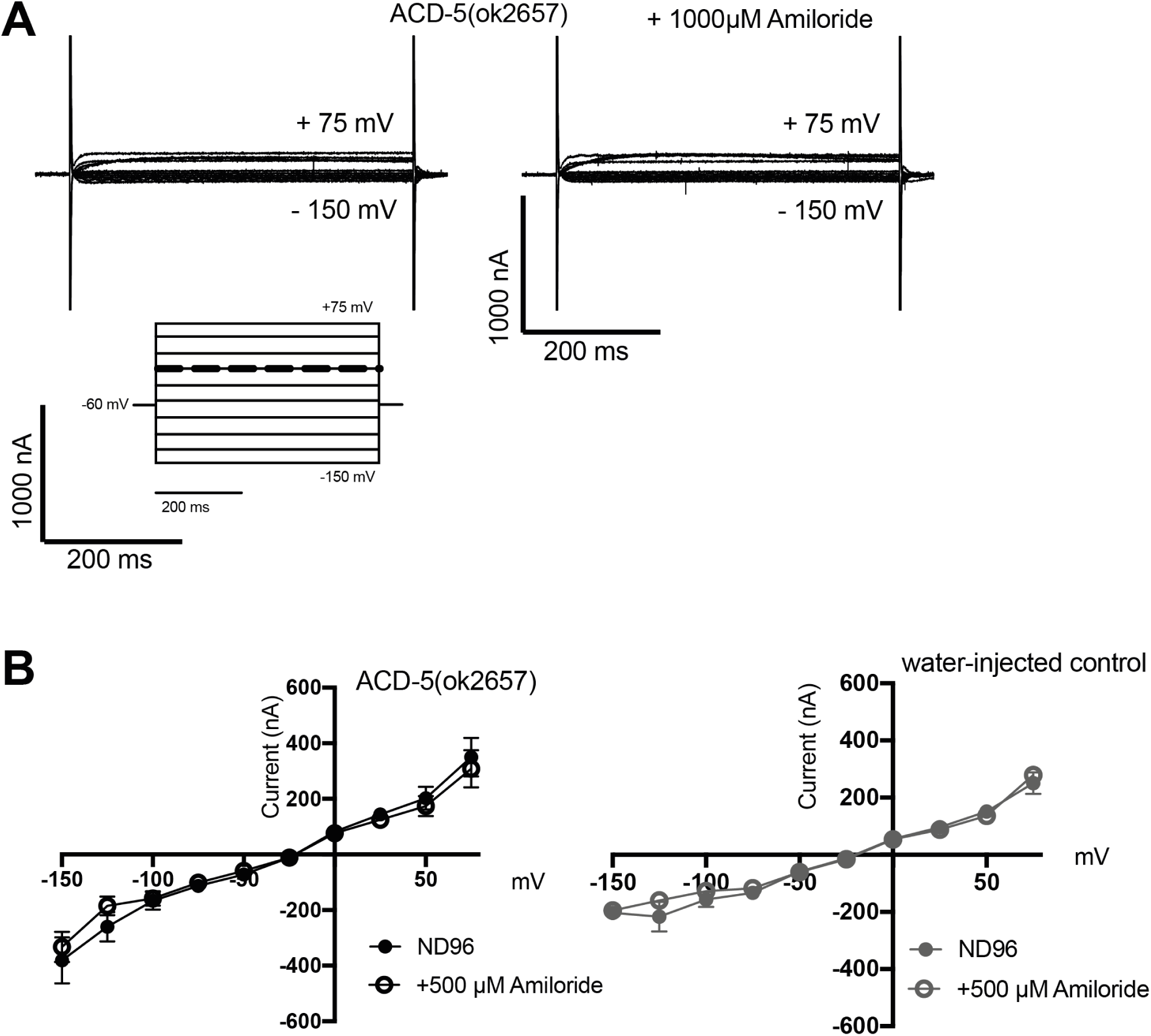
Characterization of ACD-5 channels carrying the *ok2657* mutation in Xenopus oocytes. (**A**) Representative transients and (**B**) current-voltage (IV) relationships of *Xenopus* oocytes injected with *acd-5(ok2657)* cRNA in the presence and absence of amiloride, and a water-injected control. N=10. Error bars represent mean and SEM.

**Figure 1-figure supplement 4.**
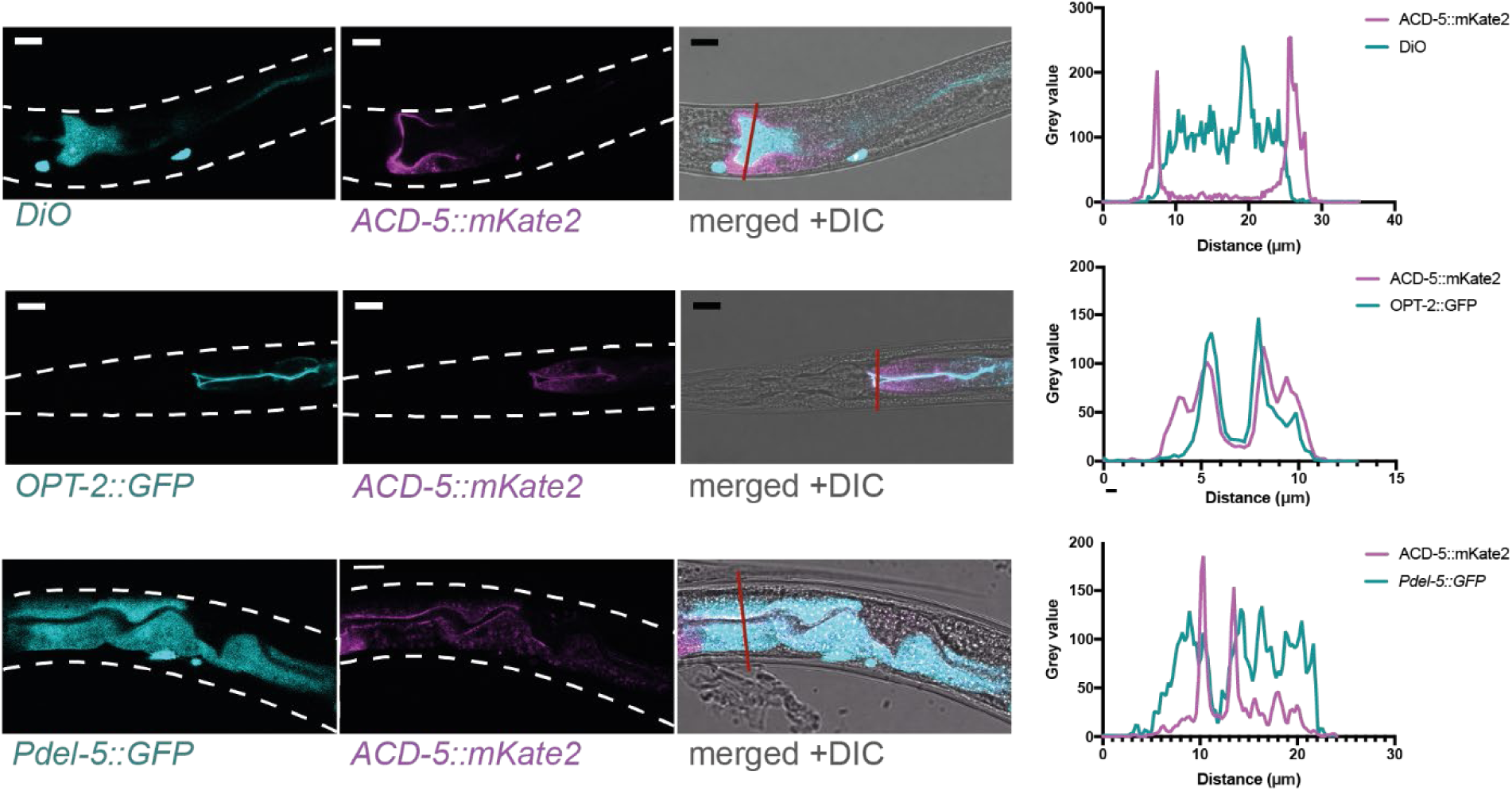
ACD-5 localizes to the apical membrane. Co-localization of ACD-5::mKate2 with luminal DiO fluorescence, an apical membrane marker *rnyEx133 [pKN114 (opt-2p:::opt-2(aa1-412)::GFP)]* and a cytoplasmic marker *ljEx1349* (*Pdel-5::GFP*) with intensity profiles. Scare bar 10 μm.

**Figure 3-figure supplement 1.**
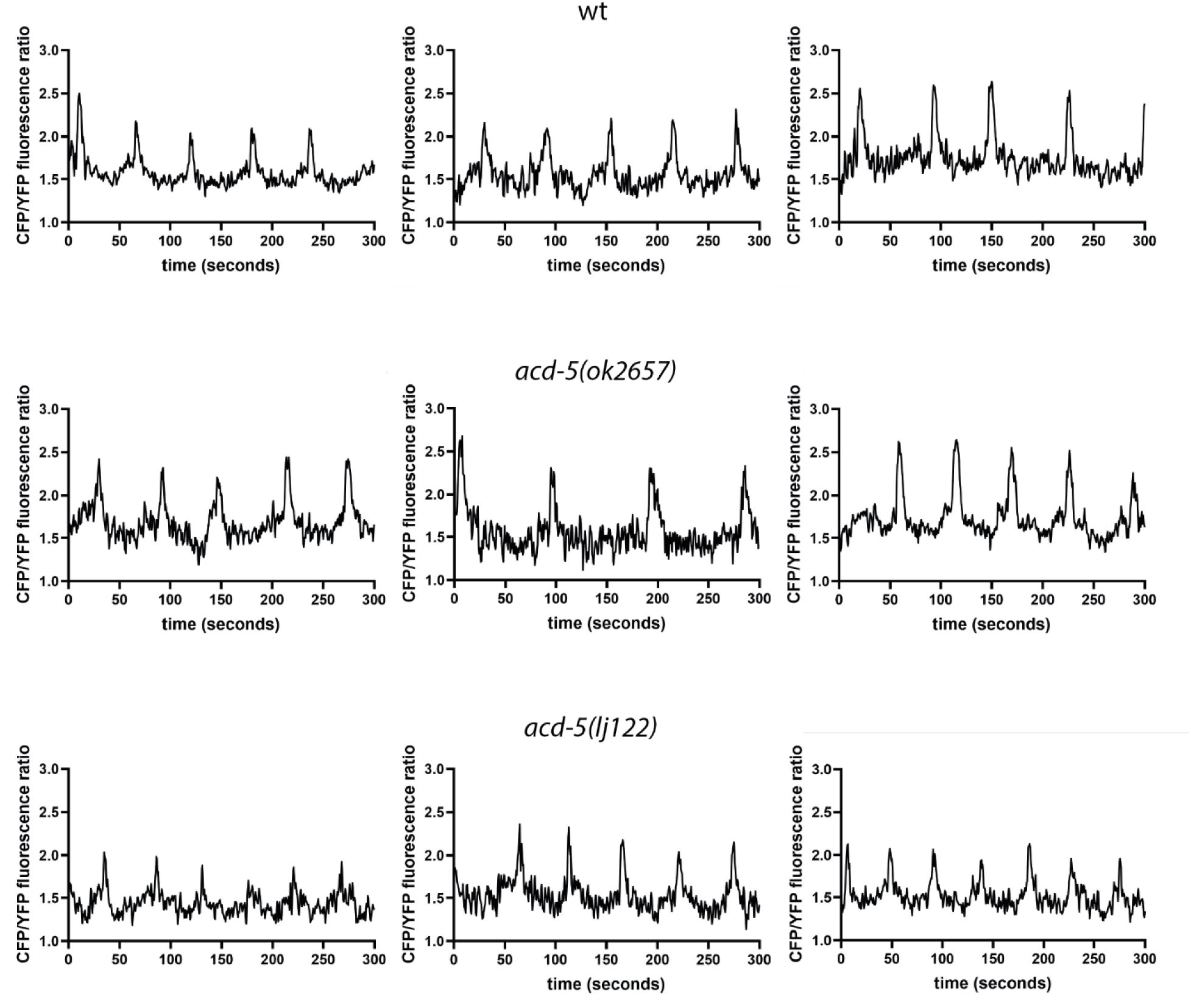
ACD-5 modifies intestinal Ca^2+^ oscillations. Ca^2+^ imaging of the intestinal cells of freely moving, defecating animals. Three representative example traces of intestinal Ca^2+^ transients of the wild type, *acd-5(ok2657)* and *acd-5(lj122)* mutants over a 5 min time period.

**Figure 3-figure supplement 2.**
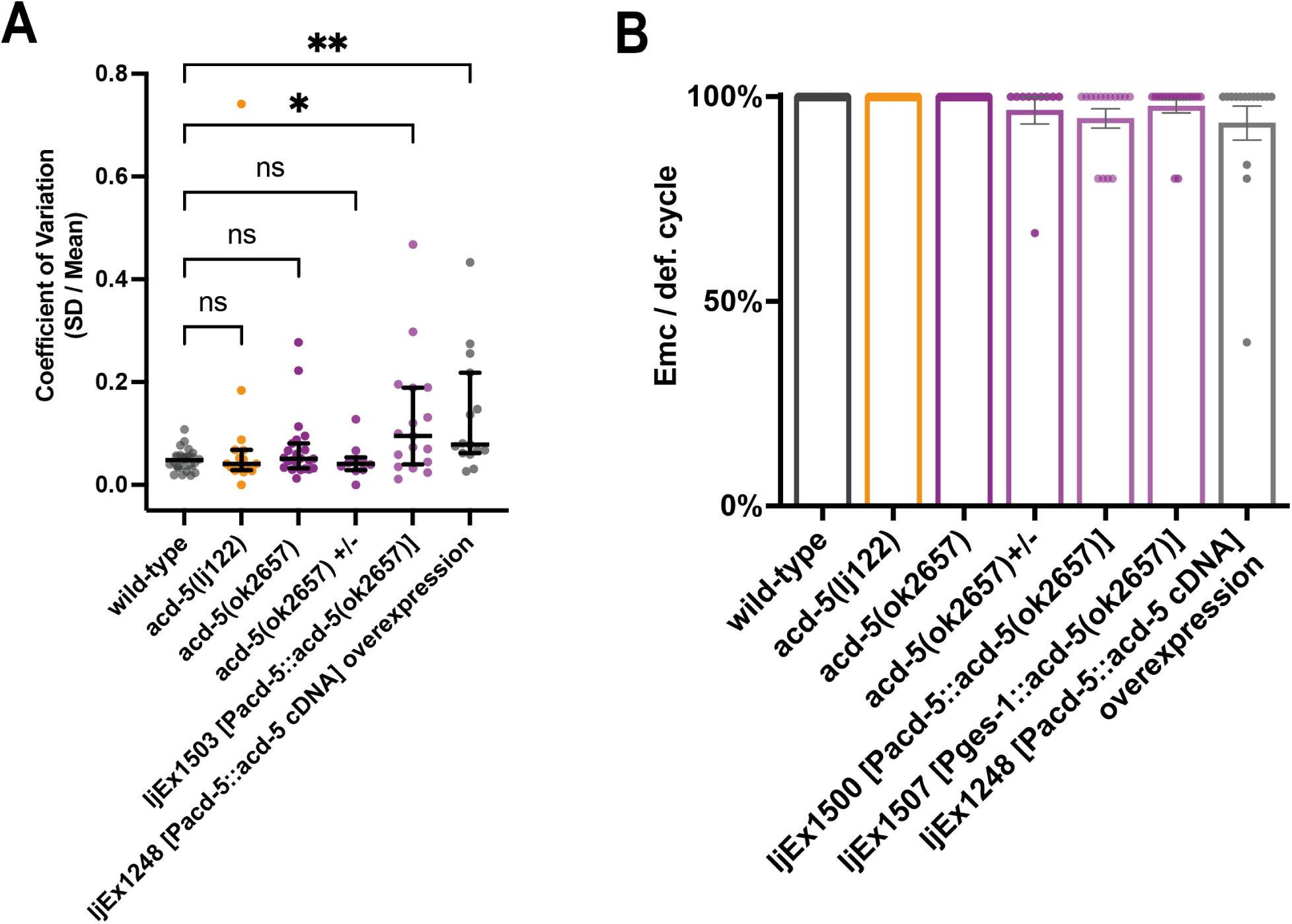
Effect of *acd-5(ok2657)* overexpression on and DMP. (**A**) Variability between DMP cycles. Error bars represent Median and IQR. Significance was assessed by a Kruskal Wallis test with Dunn’s multiple comparison test (\**p*=0.0466; \*\**p*=0.0093). (**B**) Quantification of EMCs. Error bars represent Mean and SEM. N > 10, 5 cycles each animal.

**Figure 3-figure supplement 3.**
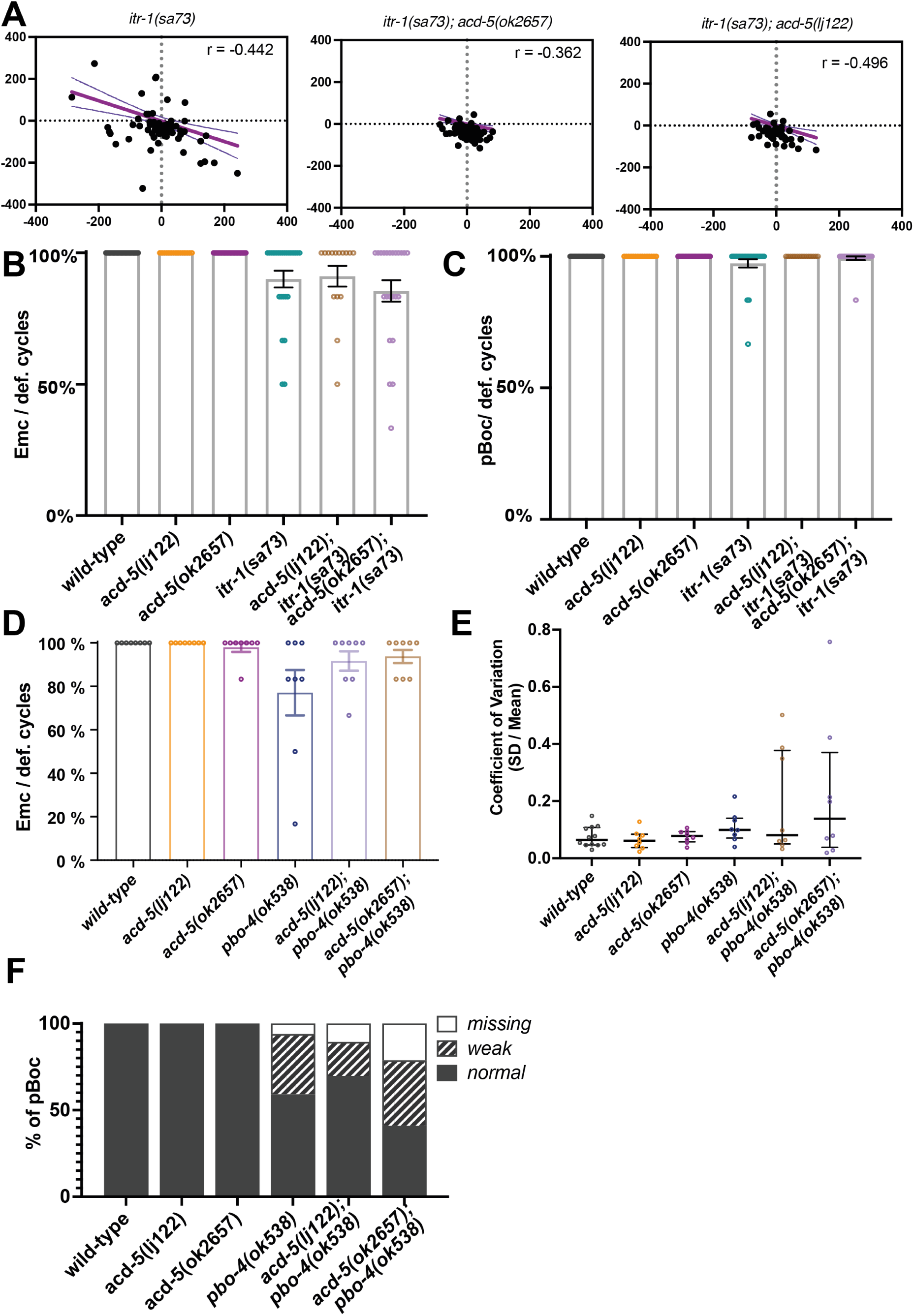
*acd-5* alleles alter the defecation phenotype of *itr-1* and *pbo-4* loss of function animals. (**A**) Analysis of the defecation cycle in *itr-1(sa73)* single and double mutants with *acd-5* reveals compensatory behavior in the oscillator. Scatter plot of the change in interval duration in one cycle pair compared to the subsequent cycle pair for *itr-1(sa73)* mutants. T(n) is the period of the cycle n. Linear regression (Pearson’s *r*) was used to derive the line, which is shown with 95% confidence limits (dashed lines). (**B**-**C**) Quantification of EMCs and pBocs executed by *itr-1/acd-5* single and double mutants. Error bars represent Mean and SEM. (**D**) Quantification of EMCs of *pbo-4* and *acd-5* single and double mutants. Error bars represent Mean and SEM. (**E**) Variability between the cycles of *pbo-4* and *acd-5* single and double mutants. Error bars represent median and IQR. (**F**) Quantification of pBoc strength of *pbo-4* and *acd-5* single and double mutants. N=15 for each condition, 5 cycles each.

**Figure 4-figure supplement 1.**
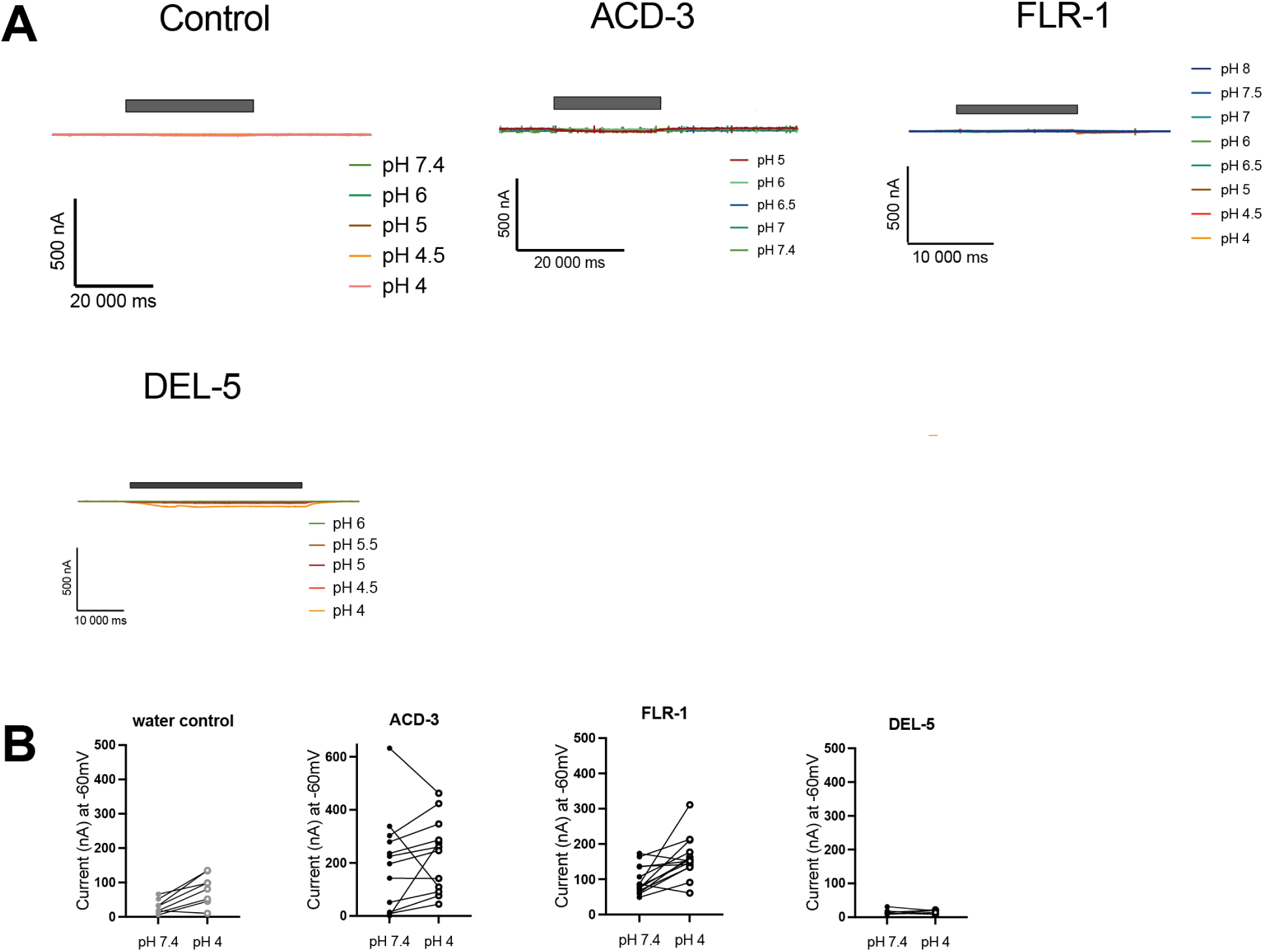
Characterization of ACD-3, DEL-5 and FLR-1 homomeric channel acid-sensing properties. (**A**) Representative traces of *Xenopus* oocytes injected with *acd-3*, *del-5* or *flr-1* cRNA (500 ng/µL) or water. (**B**) Actual currents of *Xenopus* oocytes injected with nuclease free water (control), *acd-3*, *del-5* or *flr-1* cRNA when perfused with pH 7.4 vs pH 4 (each individual oocyte is connected by a line). N = 8, 13, 14, 6 respectively.

**Figure 4-figure supplement 2.**
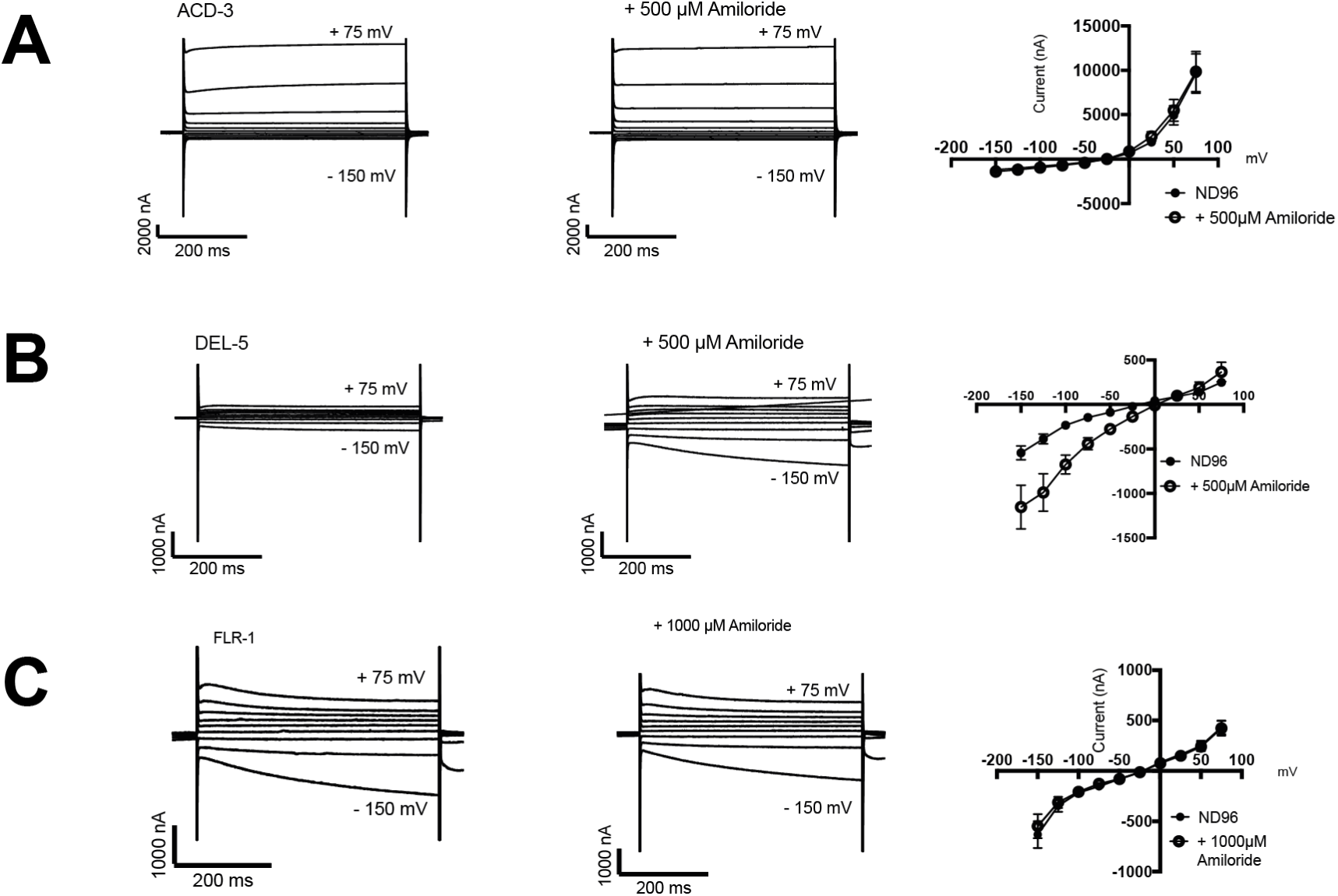
Characterization of ACD-3, DEL-5 and FLR-1 homomeric channel amiloride-sensitivity. (**A-C**) Representative traces and current-voltage relationships when *Xenopus* oocytes injected with *acd-3*, *del-5* or *flr-1* cRNA were perfused with a neutral solution (ND96, pH 7.4) in the presence and absence of 500 μM or 1000 μM amiloride. N = 5, 6, 8 respectively. Error bars represent mean and SEM.

**Figure 4-figure supplement 3.**
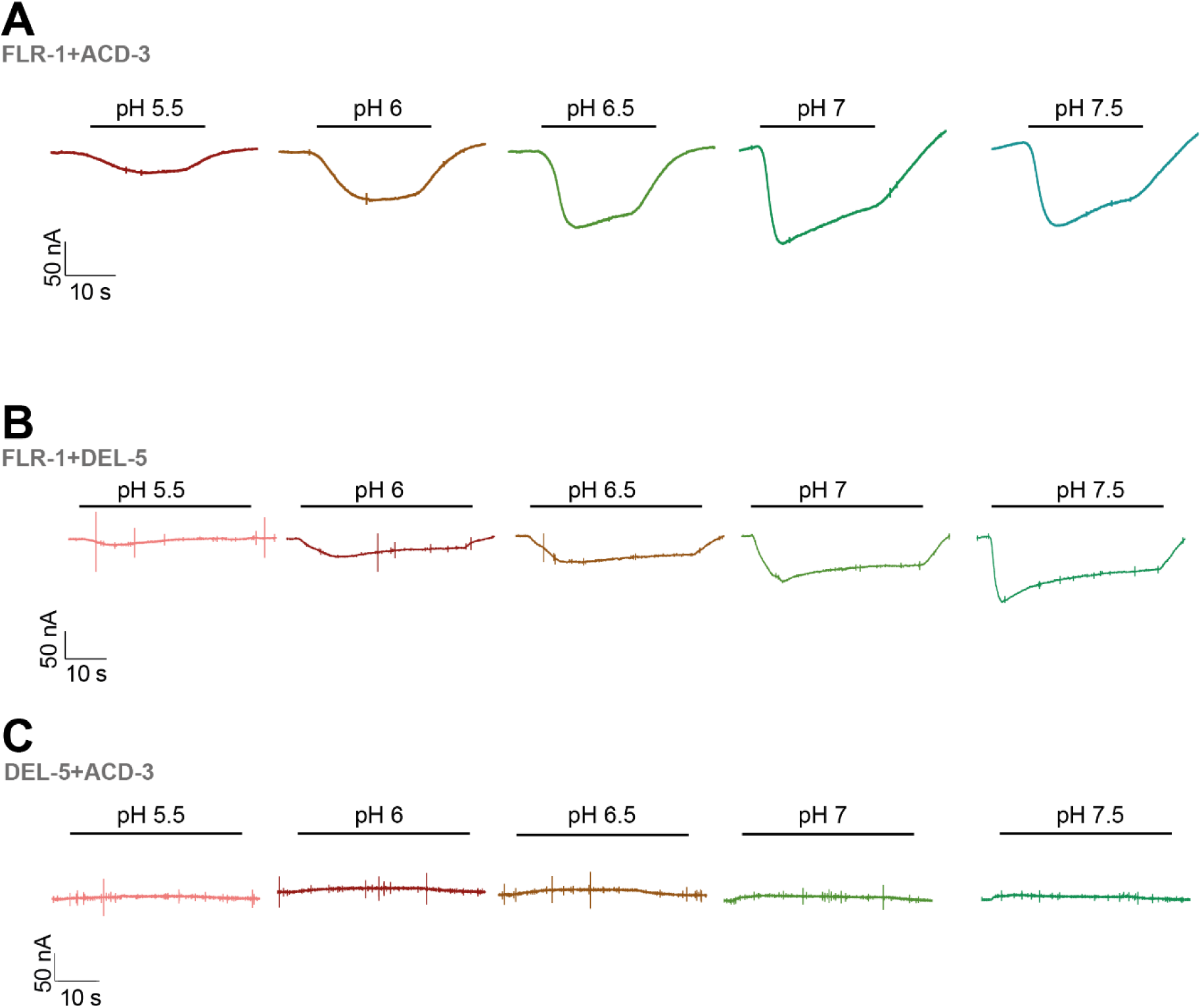
FLR-1 can form a pH-sensitive channel with ACD-3 or DEL-5. (**A-C**) Representative traces of *Xenopus* oocytes injected with *flr-1* cRNA with *acd-3* or *del-5* cRNA or *acd-3* and *del-5* cRNA (250 ng/µL for each construct). Black bar represents time of perfusion with the indicated pH. Currents were recorded at a holding potential of -60mV and traces are baseline-subtracted and drift-corrected using the Roboocyte^2+^ (Multichannels) software.

**Figure 5-figure supplement 1.**
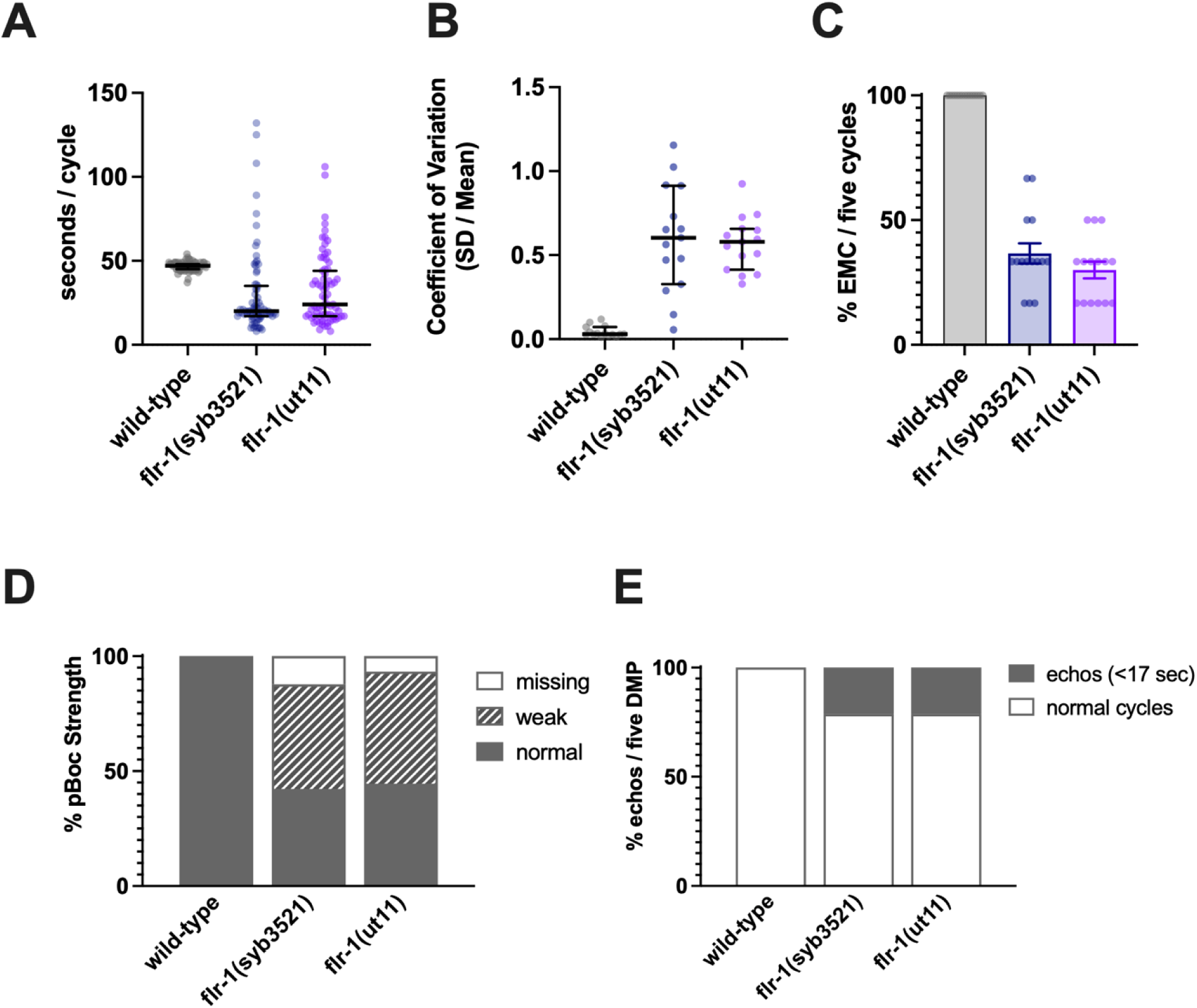
*flr-1(ut11) and flr-1(syb3521)* mutants show similar phenotypes. Comparison of *flr-1(ut110* and *flr-1(syb3521)* reveals similar defecation defects: Increased variability of cycle length, with a mix of very short and long cycles; missed EMCs and missed or weak pBocs. (**A**) Cycle length (median and IQR). (**B**) Coefficient of Variation (CV; median and IQR). (**C**) percentage of missed EMCs (Mean and SEM). (**D**) percentage of pBocs displaying the characteristics indicated. (E) percentage of cycles displaying the characteristics indicated. N = 15 animals, 5 cycles scored for each animal.

**Figure 5-figure supplement 2.**
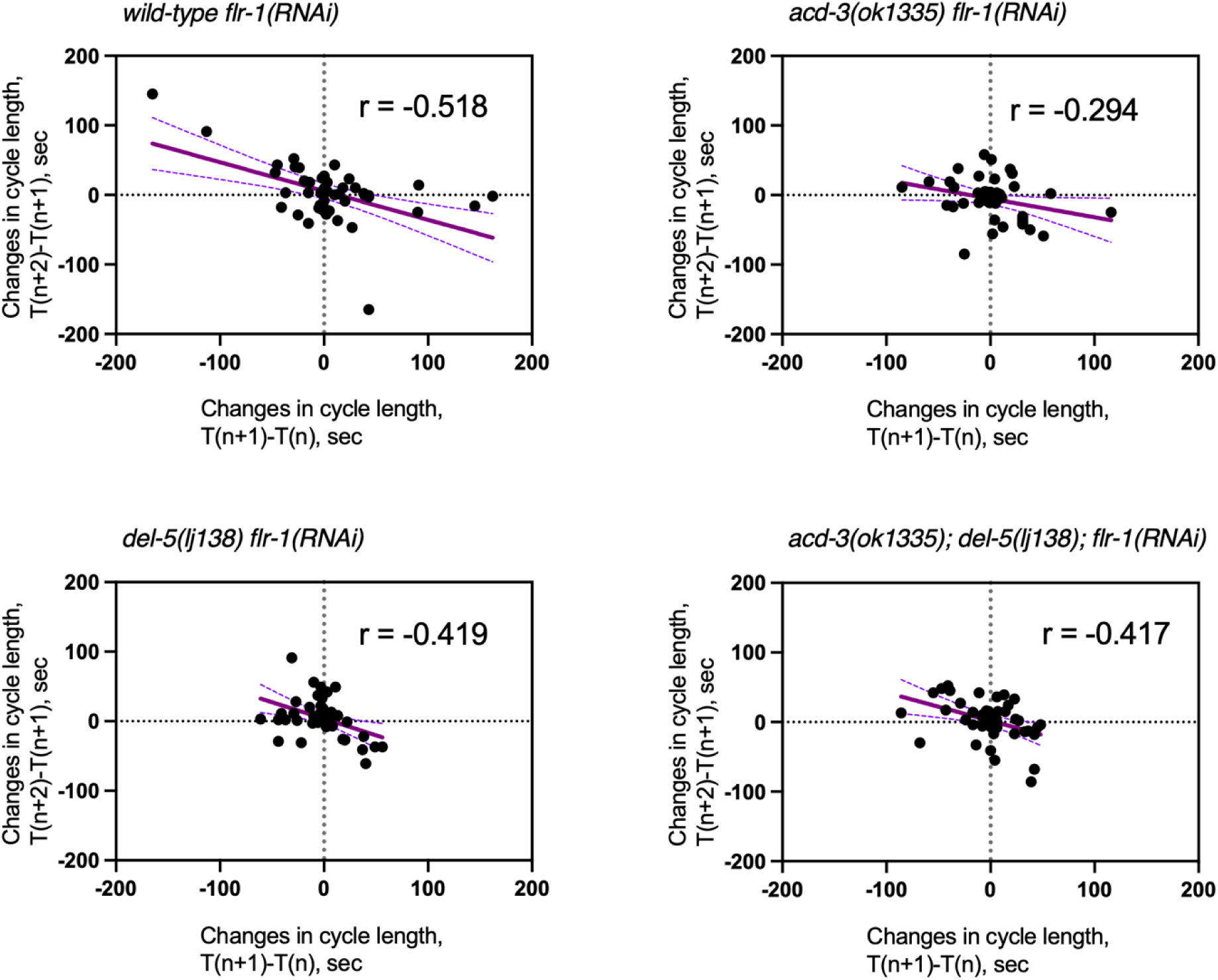
Analysis of the defecation cycle of single and double mutants on *flr-1(RNAi)* reveals compensatory behavior in the oscillator. Scatter plot of the change in period time in one cycle pair compared to the subsequent cycle pair for mutants on *flr-1(RNAi)*. T(n) is the period of the cycle n. Linear regression (Person’s *r*) was used to derive the line which, is shown with 95% confidence limits (dashed lines). N=13-15 for each genotype, 5 cycles scored for each animal.

**Figure 6-figure supplement 1.**
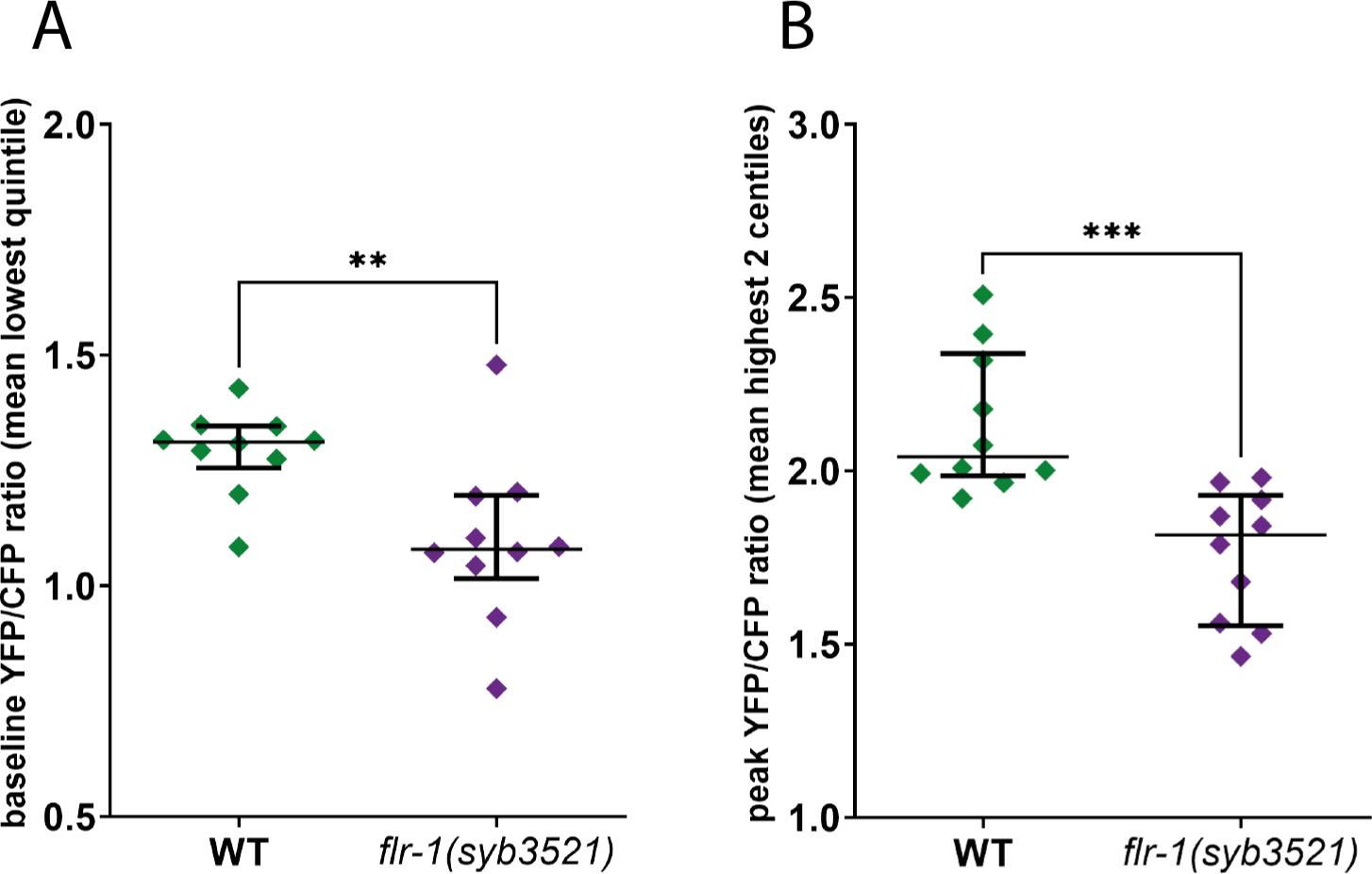
FLR-1 is required for rhythmic intestinal Ca^2+^ oscillations. Ca^2+^ imaging of the intestinal cells of freely moving, defecating animals. **(A)** *flr-1(syb3521)* animals exhibit reduced baseline Ca^2+^ levels. Scatter plot of mean YFP/CFP fluorescence ratio, for the frames corresponding to the lowest quintile, as a measure of baseline ratio, \*\**p* = 0.0076, unpaired t-test. **(B)** *flr-1(syb3521)* animals exhibit reduced maximal Ca^2+^ levels. Scatter plot of mean YFP/CFP fluorescence ratio, for the frames corresponding to the 2 highest centiles, as a measure of maximal ratio, \*\*\**p = 0.0005*, unpaired t-test. Each data point is the mean for the relevant frames, for an individual recording; error bars are SEM.

**Figure 7-figure supplement 1.**
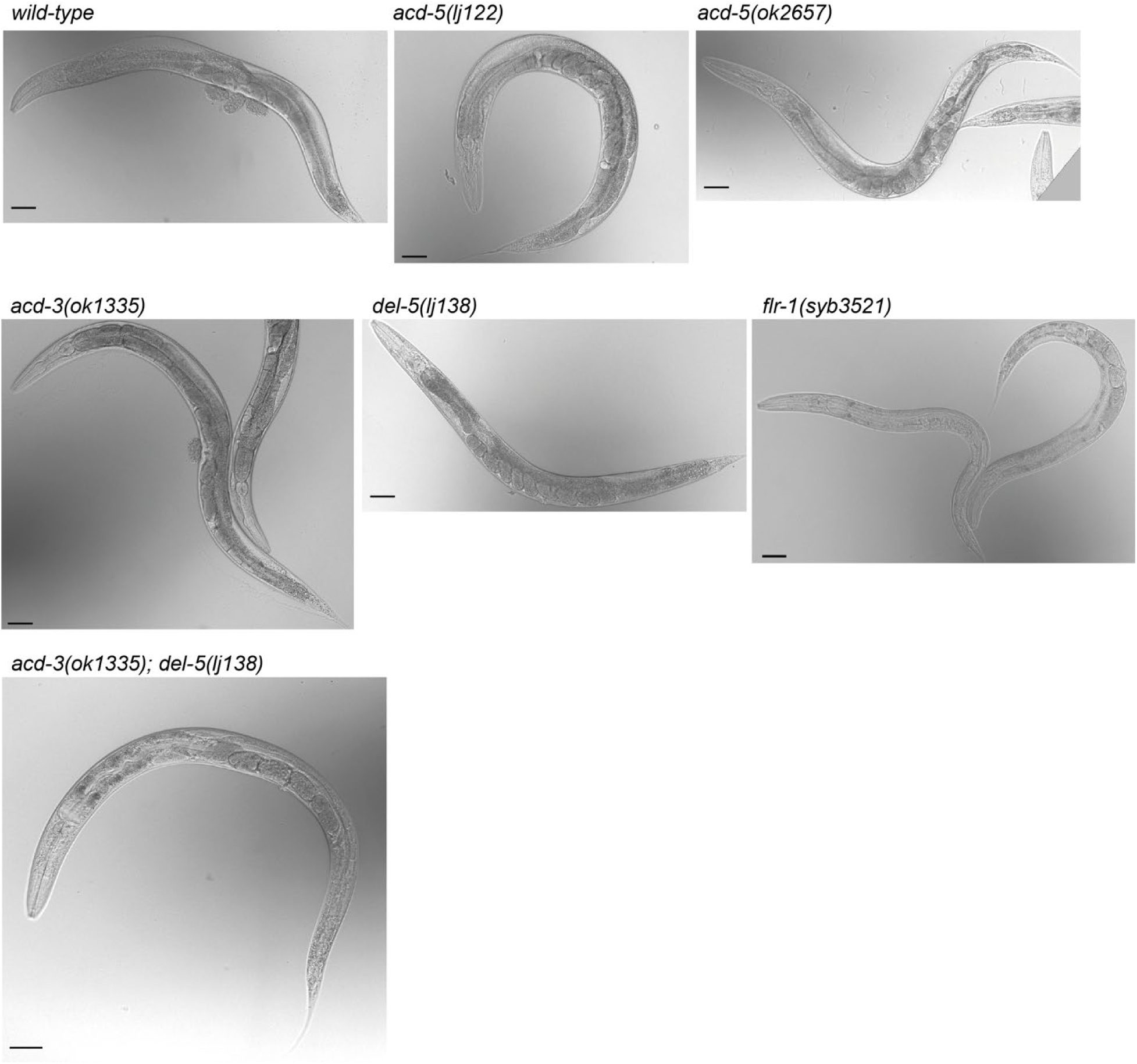
*flr-1* and *acd-3/del-5* deficient animals exhibit a caloric restriction phenotype. Single and double mutant day-1 adults for the DEG/ENaCs expressed in the intestine. Single mutants for *acd-5, acd-3 or del-5* show a wild-type body shape, while *flr-1* single or *acd-3/del-5* double mutants are thin with few eggs. Scale bars are 50 μm.

